# Regional Diversity and Leaf Microbiome Interactions of the Fungal Maize Pathogen *Exserohilum turcicum* in Switzerland: A Metagenomic Analysis

**DOI:** 10.1101/2024.04.18.590055

**Authors:** Mireia Vidal-Villarejo, Bianca Dößelmann, Benedikt Kogler, Michael Hammerschmidt, Barbara Oppliger, Hans Oppliger, Karl Schmid

## Abstract

The spread and adaptation of fungal plant pathogens in agroecosystems are facilitated by environmental homogeneity. Metagenomic sequencing of infected tissues allows to monitor eco-evolutionary dynamics and interactions betwen host, pathogen and the plant microbiome. *Exserohilum turcicum*, the causal agent of northern corn leaf blight (NCLB) in maize, is distributed in multiple clonal lineages throughout Europe. To characterize regional pathogen diversity, we conducted metagenomic DNA sequencing on 241 infected leaf samples from the highly susceptible Swiss maize landrace Rheintaler Ribelmais, collected over three years (2016-2018) from an average of 14 agricultural farms within the Swiss Rhine Valley. All major European clonal lineages of *E. turcicum* were identified. Lineages differ by their mating types which indicates potential for sexual recombination and rapid evolution of new pathogen strains, although we found no evidence of recent recombination. The associated eukaryotic and prokaryotic leaf microbiome exhibited variation in taxonomic diversity between years and locations and is likely influenced by local weather conditions. A network analysis revealed distinct clusters of eukaryotic and prokaryotic taxa that correlates with the frequency of *E. turcicum* sequencing reads, suggesting causal interactions. Notably, the yeast genus *Metschnikowia* exhibited a strongly negative correlation, supporting its known potential as biological control agent against fungal pathogens. Our findings show that metagenomic sequencing is a useful tool for analyzing the role of environmental factors and potential pathogen-microbiome interactions in shaping pathogen dynamics and evolution, suggesting their potential for effective pathogen management strategies.

## Introduction

Modern industrialized agriculture fosters adaptation and expansion of pathogens, which requires the breeding of resistant varieties and the application of chemical plant protection to maintain food security (***Savary et al., 2019***). The environmental homogeneity of modern agroecosystems, in combination with the use of resistance genes in plant breeding and plant protection chemicals, resulted in strong selection and the emergence of new virulent strains (***McDonald and Stukenbrock, 2016***). The climate change and a globalized agricultural trade additionally contribute to a rapid spread of pathogens and a more rapid pathogen evolution within agricultural systems because plant pathogens and pests spread into new geographic regions (***Chaloner et al., 2021***).

Strategic control and mitigation of pathogen proliferation require efficient disease monitoring in agricultural landscapes. Such a monitoring may employ phenotype-based diagnosis using a differentiation panel of host genotypes, each possessing unique resistance genes, or diagnostic assays using molecular markers that recognize pathogen genetic groups or races (e.g., ***Szabo et al., 2022***). These methods provide information for spatial and temporal models of plant pathogen dynamics, and elucidate factors influencing epidemiological trends. However, methods based inoculation for establishing pathogen isolates and race monitoring through a differentiation panel, are laborious, expensive, and often provide limited data. Novel strategies incorporate aerial surveillance using hyperspectral analysis (***Zhang et al., 2019***), and DNA sequencing of infected plant tissues, followed by bioinformatic and population genetic analyses (***Hubbard et al., 2015***). Genomic information from infected tissues allows to analyze pathogen epidemiology and demographic history, characterize multiple infections by different strains, and identify new strains arising from mutation, sexual recombination, or introgression from other regions. Additionally, metagenomic analysis of sequencing data from infected tissues provides valuable data on co-infecting eukaryotic and prokaryotic organisms (***Regalado et al., 2020***). Such data contribute to an understanding of eco-evolutionary interactions between pathogens and associated microbes, providing a foundation for future breeding and disease control strategies.

The fungal pathogen *Exserohilum turcicum* (telomorph *Setosphaeria turcica*) is a major worldwide maize pathogen causing northern corn leaf blight (NCLB) (***Galiano-Carneiro and Miedaner, 2017***). NCLB is a significant maize disease in Central Europe due to its extensive cultivation and highly homogenous agricultural landscape. Rapid breeding progress and expansion of maize cultivation areas have made it a dominant crop in European agriculture, impacting the epidemiology and evolution of *E. turcicum*. The pathogen rapidly expanded throughout Southern and Central Europe since the late 19th century, leading to yield reduction until breeding programs selected for resistance (***Galiano-Carneiro and Miedaner, 2017***). A large-scale monitoring of *E. turcicum* races using different maize genotypes with monogenic resistance genes revealed diverse races with a distinct geographic distribution (***Hanekamp, 2016***). In a previous study, we characterized the geographic distribution of *E. turcicum* genetic variation using whole genome sequencing and found independent introgressions into Europe and varied distribution patterns of genetic lineages (***Vidal-Villarejo et al., 2023***). However, the resolution of pathogen genetic diversity dynamics is limited when isolates are sampled at a single point in time, particularly at smaller temporal and geographic scales.

Here we describe metagenomic sequencing of naturally infected leaves of a highly susceptible European maize landrace to monitor small-scale spatial and temporal patterns of *E. turcicum* genetic diversity. We first conducted a pilot experiment to implement appropriate protocols and analysis pipelines by collecting leaf samples from a small subset of commercial maize hybrid varieties. Subsequently, we analyzed over 200 leaf samples taken from the highly susceptible traditional variety, Rheintaler Ribelmais, which had been collected over three consecutive years (2016-2018) from agricultural production fields located within the Rhine Valley in Switzerland. Our objectives included characterizing the genetic lineages of *E. turcicum* present within the Rhine Valley and comparing them with those found throughout Europe. Using the sequencing data we tested whether levels of genetic diversity of this pathogen in the Swiss Rhine valley is lower than throughout Europe and whether there is a relationship between spatio-temporal patterns of phyllobiome and pathogen diversity that reflect causal interactions.

## Results

### Pilot experiment for method development

We first conducted a pilot study to develop DNA extraction and sequencing protocols and determine sequencing parameters that result in amounts of pathogen DNA. The experiment included collecting a total of nine metagenomic leaf samples from three different plants of each of three commercial maize hybrid varieties (abbreviated as R, G and T) of a private breeding company, which had a high level of resistance to *E. turcicum* and were naturally infected in a field in Salez/Switzerland. After library construction and sequencing (see Materials and Methods), we obtained 55.4 GB raw short read sequencing data. The percentage of reads in these nine metagenomic samples that mapped to the *E. turcicum* reference genome ranged from 7.72% to 22.92%, with estimated coverages from 9.44X to 32.4X (see Supplementary Dataset S1). The data also include two *E. turcicum* isolates from Rheinau and Hombrechtikon, Switzerland (13.4 GB raw data) used as a control of the metagenomic procedures for the identification of *E. turcicum* lineages. Summary statistics of metagenomic samples indicate that approximately 10% of sequence reads from infected leaves map to the *E. turcicum* reference genome. Therefore, to obtain about 20X coverage of *E. turcicum* in a single infected leaf sample, 8.6 GB of raw output per sample is needed.

### Regional sampling and sequencing of infected maize leaves

We then collected leaves from the susceptible traditional cultivar Rheintaler Ribelmais, which were naturally infected with *E. turcicum*, from fields across the Rhine Valley in Switzerland over three consecutive years (2016, 2017, and 2018; Fig. 1).

**Figure 1.**
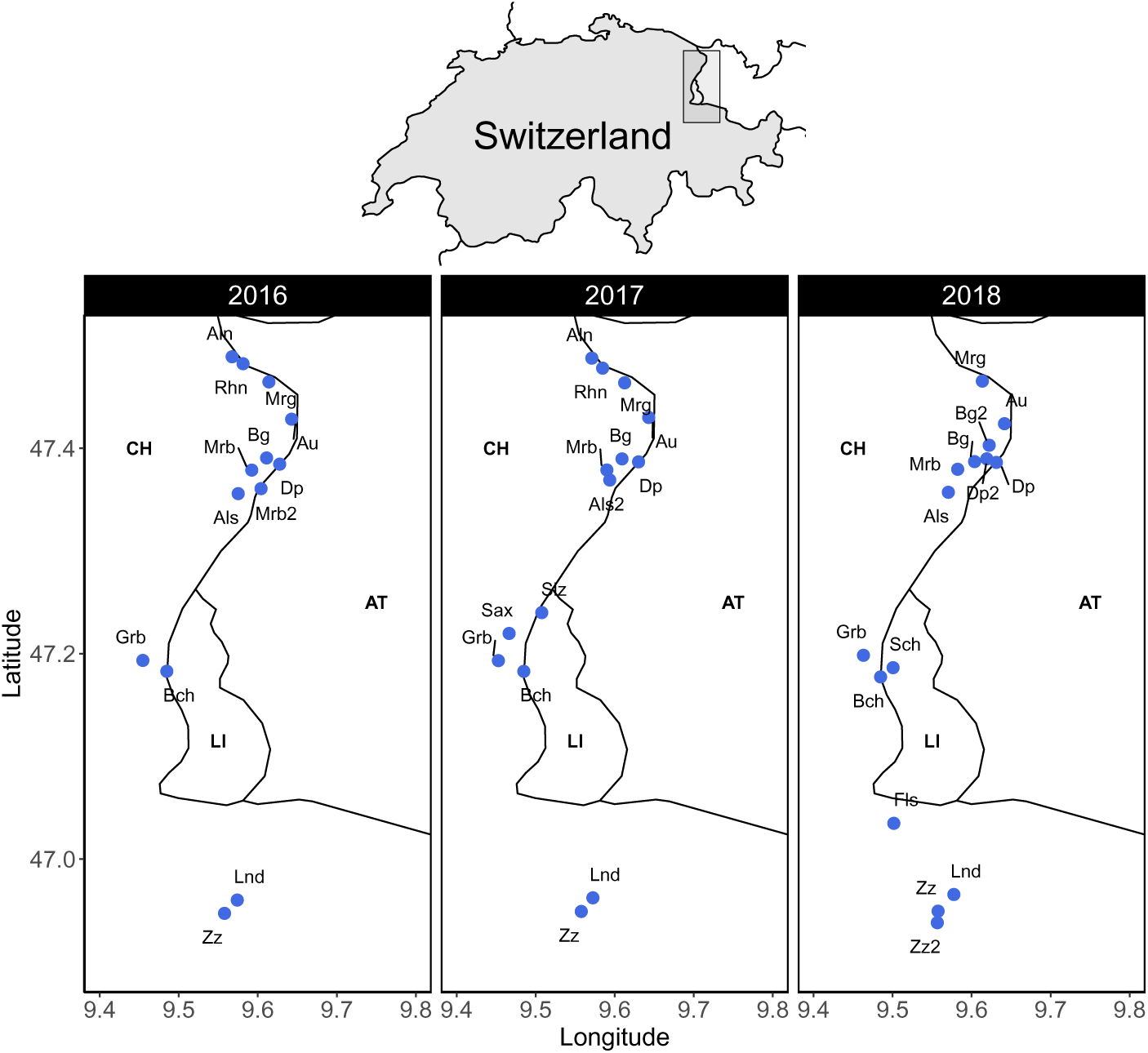
Geographic location of agricultural fields in the Upper Rhine Valley of Switzerland (CH) along the border to Austria (AT) and Liechtenstein (LI) from which infected maize leaf samples were collected over three years. In text, abbreviation of the collection sites (see Supplementary Dataset S1 for complete location names).

Each year, we collected infected leaves from an average of 14 different locations, resulting in a total of 21 different agricultural fields. Nine fields were sampled in all three years, three fields in two years, and nine fields only once. For each site and year, we separately sequenced one leaf each from six to eight different plants per field, resulting in a total of 247 metagenomic sequencing samples and 2,547.34 GB of raw data. We discarded six samples with a low number of sequencing reads (<1,000 reads). Supplementary Dataset S1 provides detailed information about the sampling location and a summary of the sequencing data.

We first estimated the approximate proportion of *E. turcicum* and maize reads in the metagenomic sequencing data by mapping all trimmed reads to *E. turcicum* St28A and maize AGPv4 reference genomes, respectively. Across all 241 samples, a median of 6.35% of reads mapped to *E. turcicum* and 47.32% to maize, although there were differences between years (Fig. 2 and Supplementary Fig. S1). Samples from 2018 had higher proportion of reads mapping to maize than samples from 2016 and 2017, whereas the proportion of reads mapping to *E. turcicum* was similar. On average, the 2016 samples showed the highest percentage of reads mapping to *E. turcicum*, especially samples from fields in Buchs (Bch), Balgach (Bg), and Grabs (Grb).

**Figure 2.**
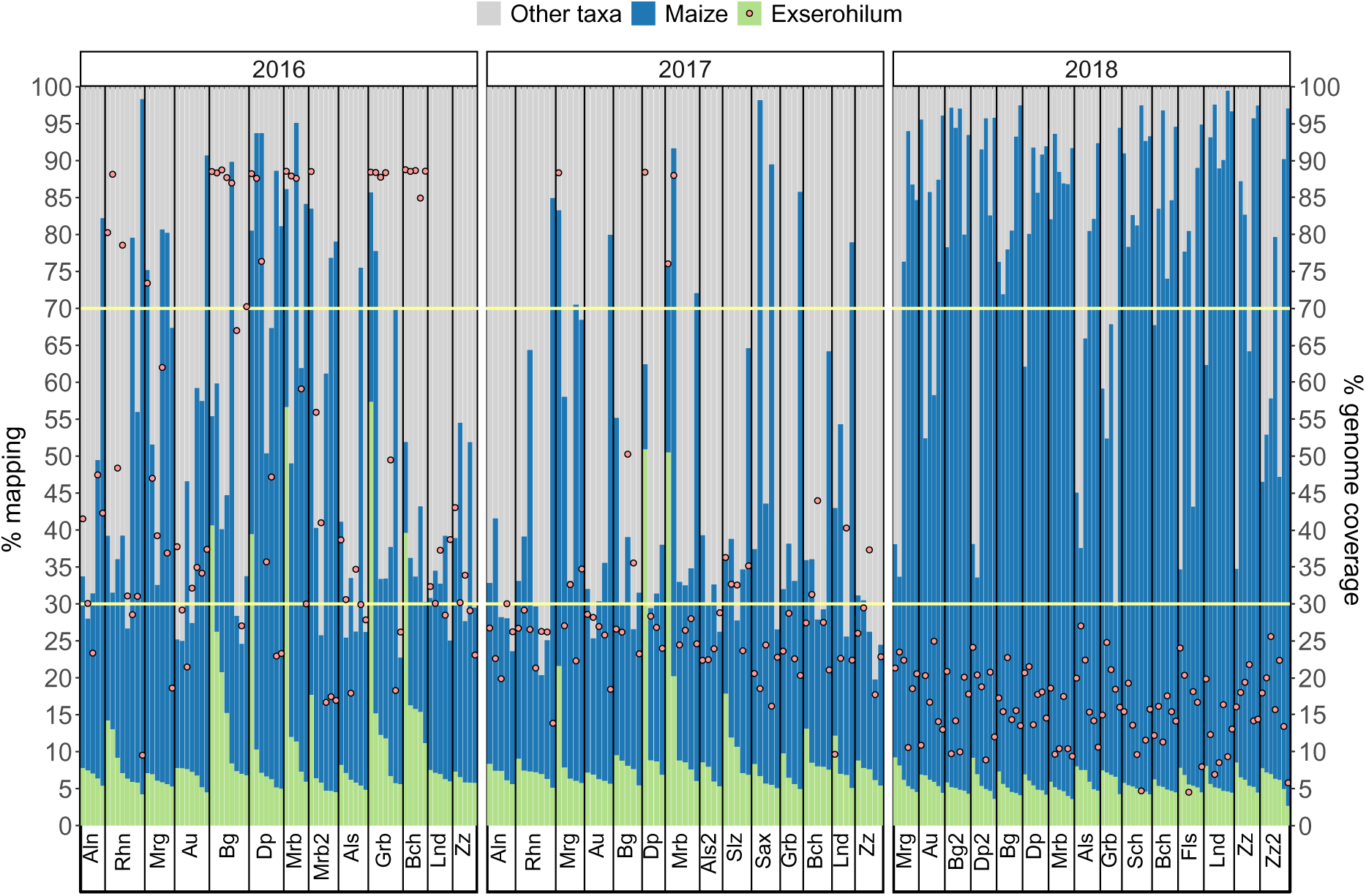
Percentage of reads mapping to *E. turcicum* and maize reference genomes (left y-axis). ‘Other taxa’ represent reads that neither map to *E. turcicum* nor maize. Red dots indicate the percentage of geonme coverage of *E. turcicum* (right y-axis). The samples are grouped by year and collection site as indicated in the x-axis. Collection sites (names abbreviated, see Supplementary Dataset S1 for the complete names) are ordered by latitude, i.e., in North to South direction and samples within location ordered by % of reads mapping to *E. turcicum*. The horizontal yellow lines highlight 70% and 30% of *E. turcicum* genome coverage.

Of the 241 sequenced samples, 30 samples covered ≥ 70% of the *E. turcicum* genome, whereas 165 samples incompletely covered (≤ 30%) the genome despite having similar percentage (5-7%) of reads mapping to *E. turcicum* (Fig. 2).

Since a direct mapping of metagenomic reads can lead to misclassification of *E. turcicum* reads due to the presence of reads from its close relatives in the reference genome, we used metagenomic classification to classify reads into different taxa first and then map *Exserohilum* reads to the reference genome.

### Taxonomic read classification

To identify reads originating from *E. turcicum* and to characterize the dynamics of the taxonomic diversity of infected Rheintaler Ribelmais leaves over years and field locations, we classified metagenomic sequencing reads at the genus level using Kaiju (***Menzel et al., 2016***). This tool identified 384 eukaryotic and 258 bacterial genera. The ten most abundant eukaryotic genera accounted for 69.0% of the eukaryotic reads and included the genera *Epicoccum* (25.8%), *Alternaria* (16.1%), *Exserohilum* (5.6%), *Puccinia* (4.3%), and *Bipolaris* (4.3%). Although DNA was extracted from leaves with symptoms of an *E. turcicum* infection, *Exserohilum* was the dominant taxon in only a few samples from 2016 and 2017 (Fig. 3). Among bacteria, the top ten genera accounted for 74.4% of the bacterial reads and included the genera *Acinetobacter* (21.0%), *Pantoea* (11.8%), *Sphingomonas* (11.7%), *Hymeonbacter* (5.7%), and *Pseudomonas* (5.1%).

**Figure 3.**
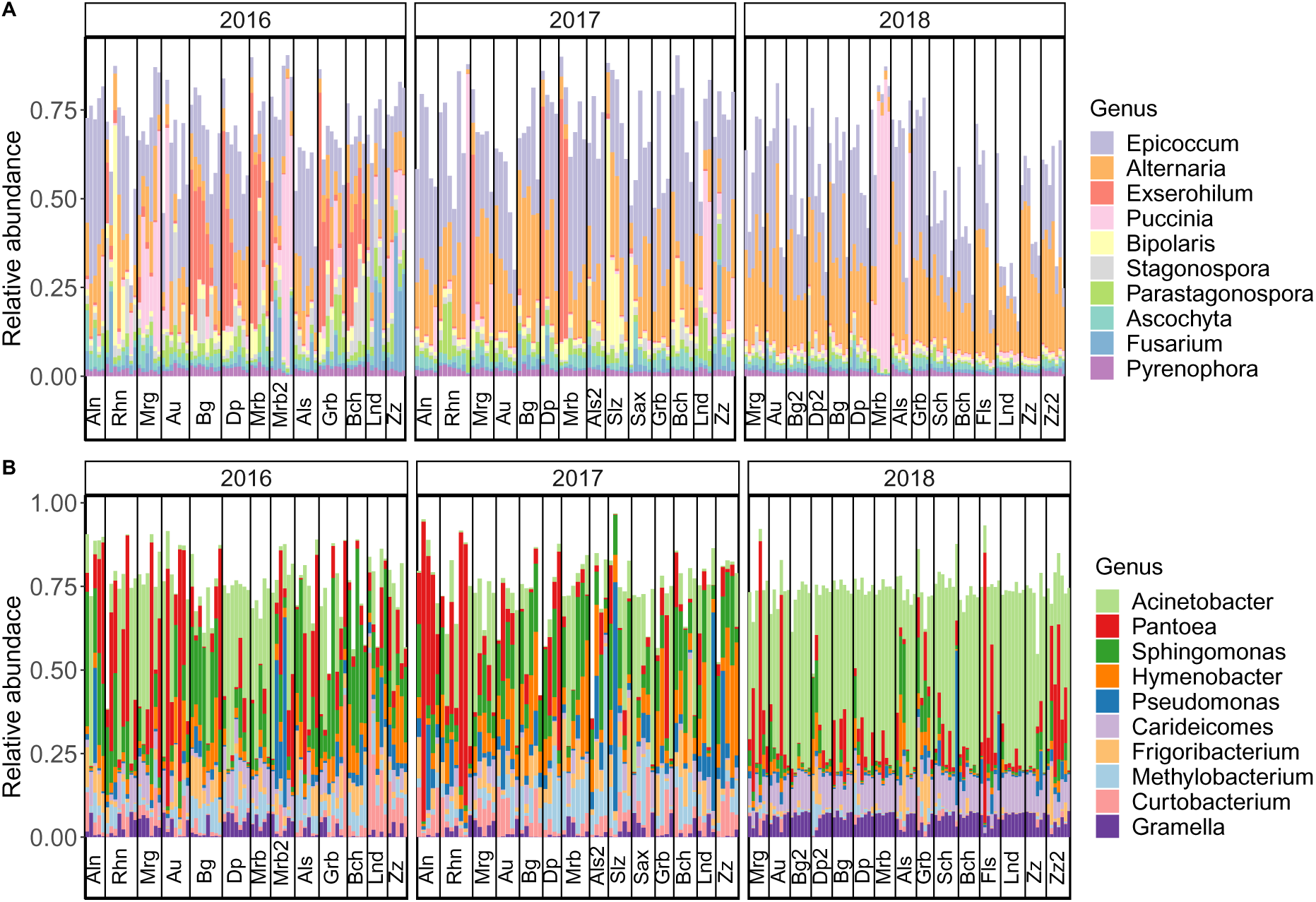
Per sample relative abundances of the global top 10 genera of A) Eukarya and B) Bacteria in the years 2016 to 2018. The samples are ordered by collection site from North to South and by percent *Exserohilum* sequence reads within sites. Collection site names are shown abbreviated, see Supplementary Dataset S1 for complete names

### Composition of taxa in the phyllobiome over space and time

We estimated the diversity of the phyllobiome using Shannon’s index, 𝐻^′^. In all three years, eukaryotic diversity was higher than bacterial diversity (Paired t-test, 𝑝 = 3.52 × 10^−09^ in 2016; 𝑝 = 0.00135 in 2017; and 𝑝 < 2.2 × 10^−16^ in 2018). In 2016 and 2017, there were no significant differences in eukaryotic and bacterial diversities (Wilcoxon test, 𝑝 > 0.01). In 2018, the bacterial diversity was lower and the eukaryotic diversity was higher compared to the previous years (Fig. 4A).

**Figure 4.**
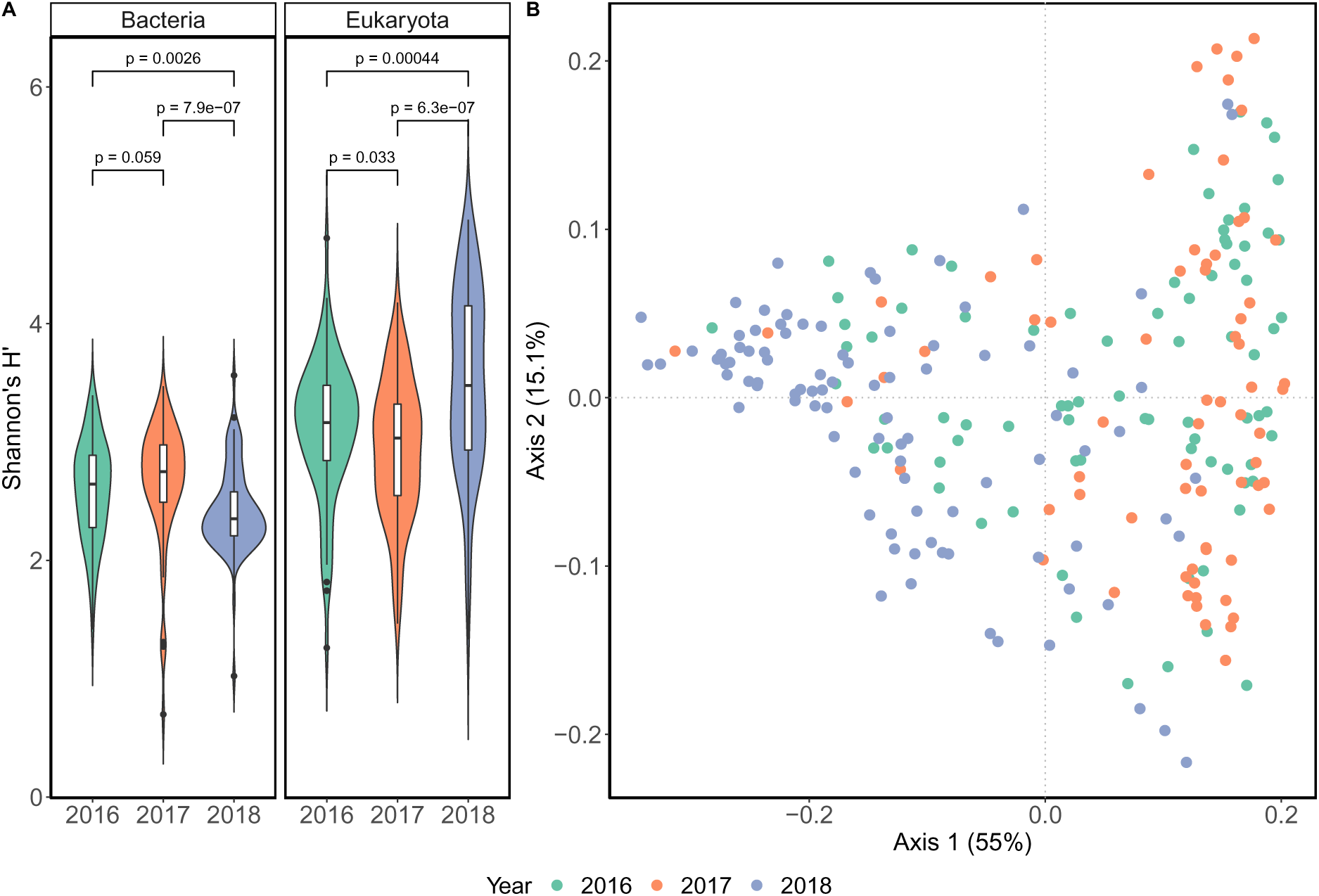
A) Shannon’s diversity index, 𝐻^′^, for the bacterial and eukaryotic phyllobiome over three consecutive years. The 𝑝 value of the Wilcoxon rank test between years is shown. Sample sizes for 2016, 2017, and 2018 were 80, 69, and 92, respectively. B) The first two principal coordinates of a PCoA analysis conducted with Bray-Curtis dissimilarities of the relative abundances of Eukaryota and Bacteria. Colors indicate the sampling year, and the percentage of variance explained by each axis is indicated in parenthesis.

A Permutational Multivariate Analysis of Variance (PERMANOVA) of phyllobiome dissimilarities through Bray-Curtis distances indicates an association (𝑝 < 0.001) between phyllobiome composition and sampling year (𝑅^2^ = 0.242) and collection site (𝑅^2^ = 0.112), respectively, and a sampling year and collection site (𝑅^2^ = 0.078, 𝑝 < 0.01; Table 1). There is no association (𝑝 > 0.05) with phyllobiome dissimilarities and latitude nor longitude. Similar results were obtained when eukaryotic and bacterial reads were analyzed separately (Supplementary Table S1 and Supplementary Fig. S2). The previous analysis of variance does not tell whether the differences observed in years are due to differences between all three years or between specific years. We therefore conducted a pairwise PERMANOVA between years of sampling. This analysis reveals that the taxonomic composition in 2018 differs the most among the three years (Table 1). The pairwise comparison of taxonomic diversity between 2016 and 2017 shows little variation explained by the year (𝑅^2^ = 0.031, 𝑝 = 0.0014). In contrast, pairwise comparisons between 2016 and 2018 (𝑅^2^ = 0.227, 𝑝 = 1𝑒 − 05) and between 2017 and 2018 (𝑅^2^ = 0.279, 𝑝 = 1𝑒 − 05) suggest greater variation. These results are consistent with the PCoA analysis, in which the first axis separates most samples from 2018 from those of 2016 and 2017, whereas the second axis does not differentiate between samples from 2016 and 2017 (Fig. 4B).

**Table 1.**
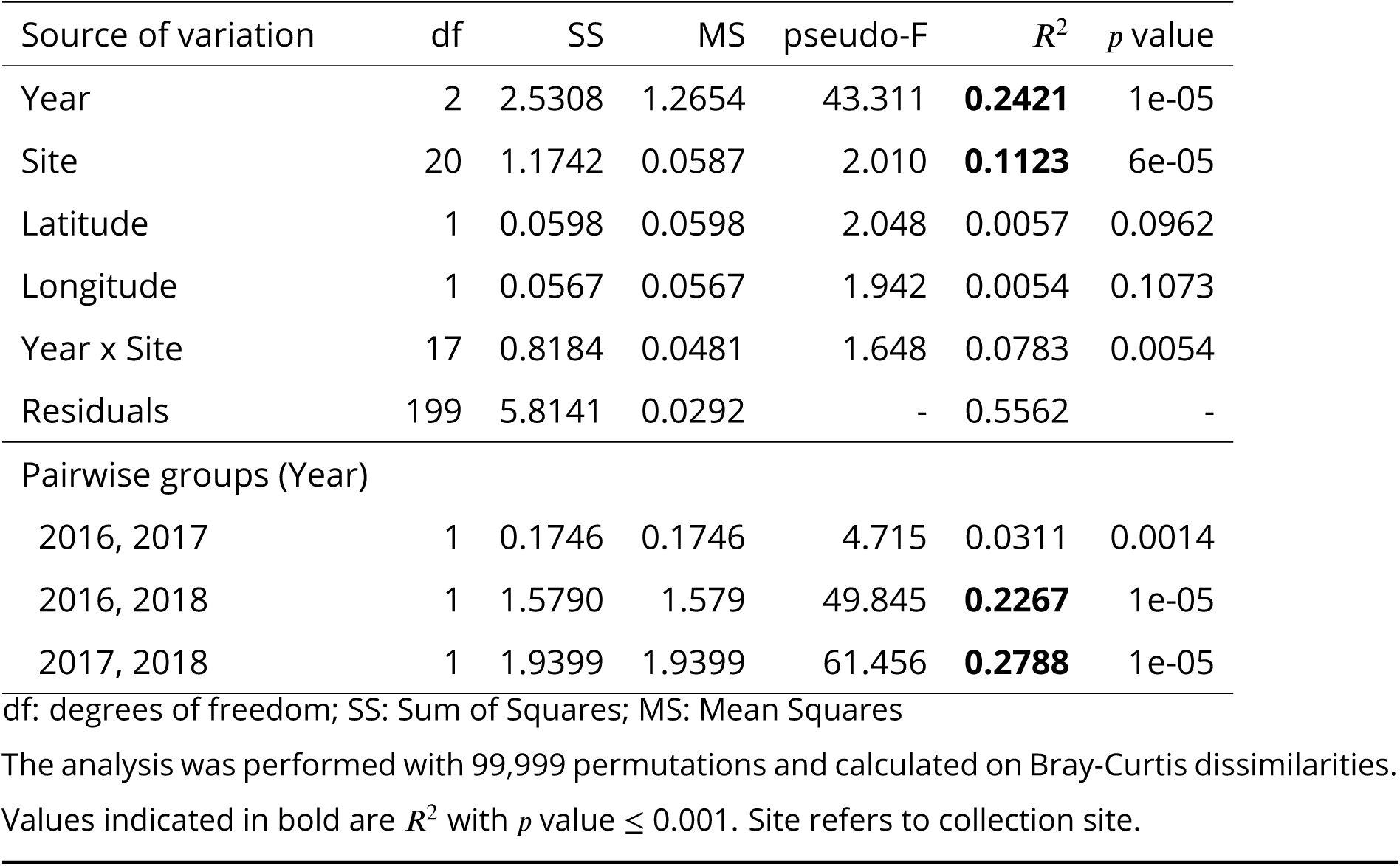
Multifactorial PERMANOVA test statistics for significance factors influencing phyllobiome dissimilarities and pairwise PERMANOVA on the year.

A permutation test of homogeneity of dispersion found that the dispersion in 2018 differed from 2017 and 2016. To validate that the differences observed in the PERMANOVA do not result from a the unequal dispersion, we performed the PERMANOVA by subsampling 20 random samples from each year 500 times, following ***Karasov et al. (2019)***. In most subsamples, the difference in dispersion is no longer significant (mean 𝑝 = 0.19), while PERMANOVA remains significant (mean 𝑅^2^ = 0.25 and mean 𝑝 = 8 × 10^−05^).

When conducting separate PERMANOVA analyses for each collection site, we observed a consistent and robust association between the taxonomic composition and the year of sampling in most sites where data were available for all three years. In all analyses, the permutation test of dispersion provided no statistical support (𝑝 > 0.01). However, two specific locations, namely St. Margarethen (Mrg) and Grabs (Grb), did not show a significant association with the year of sampling (Supplementary Fig. S3).

Since the differences in phyllobiome composition between years can mainly be attributed to the year 2018, we used publicly available meteorological data from April to September to test whether differences in weather may explain this observation. The agrometeorological data for the Swiss Rheintal region (by month) over the three years show lower leaf wet duration, total precipitation, and mean relative humidity in 2018, but higher evapotranspiration and mean temperature compared to the two previous years (Wilcoxon rank test, 𝑝 value < 0.01 for all analyses, evapotranspiration between 2017 and 2018 with 𝑝 value of 0.017; Fig. 5). Leaf wet duration, evotranspiration, total precipitation, mean temperature were not different between 2016 and 2017 (Wilcoxon rank test; 𝑝 value > 0.01). These differences are not mirrored when analysing extreme values (i.e., the maximum and minimum temperature and humidity measurements) within the same monthly period (Supplementary Fig. S4). This suggests that annual variation in taxonomic composition and diversity of the phyllobiome is mainly influenced by the average weather conditions and less by days of extreme weather.

**Figure 5.**
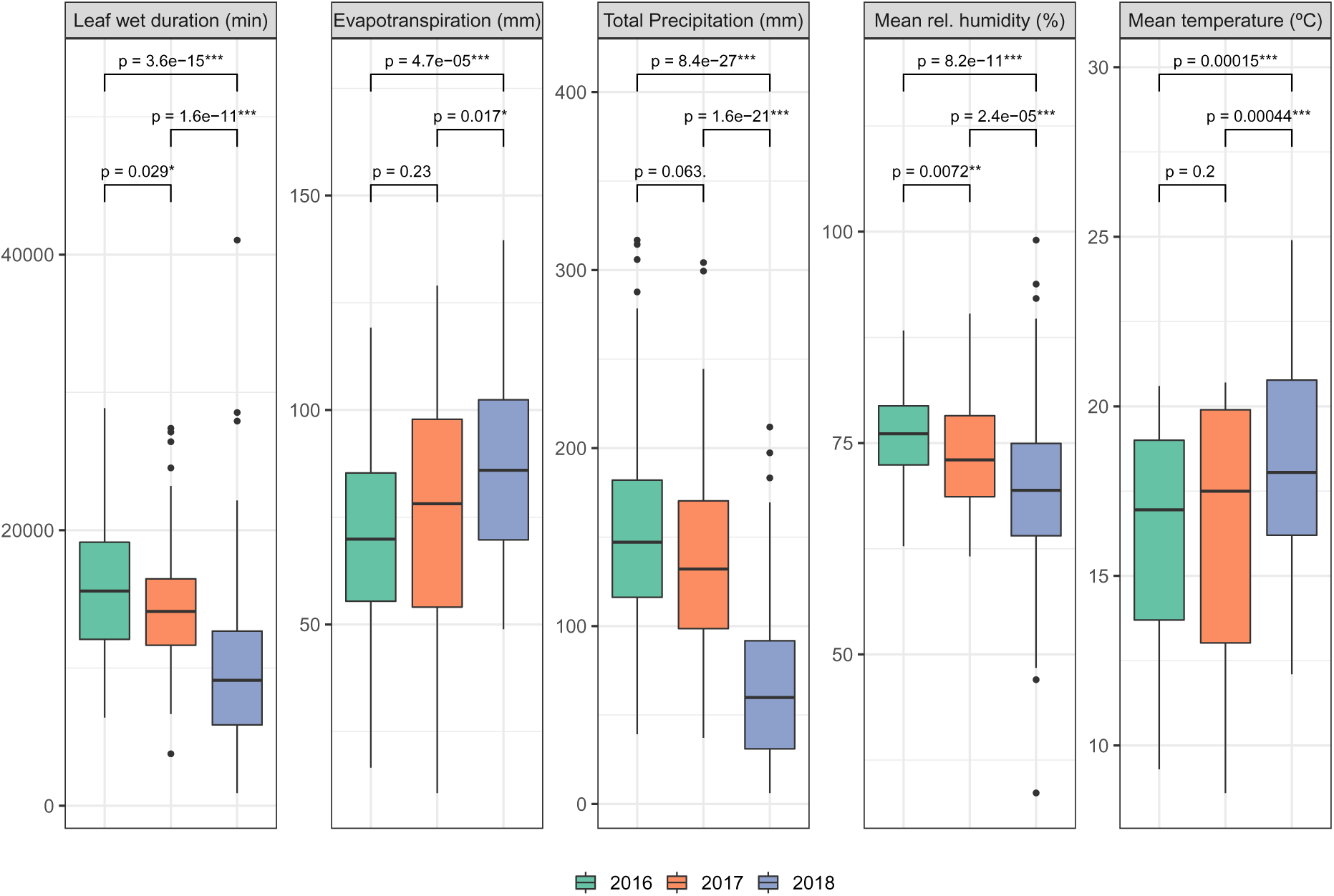
Agrometeorological monthly data from Rheintal summarized by each year, including months from April to September. p indicates 𝑝 value of Wilcoxon rank test between years.

We also investigated the variation in taxonomic composition among the three maize hybrid varieties in the pilot study. The comparison of Shannon’s index, 𝐻, across the three varieties did not yield any significant differences (Supplementary Fig. S5). A PERMANOVA analysis using the maize variety as factor did not reach statistical significance (𝑝 = 0.1524). This finding suggests a limited effect of host genetic background on microbiome diversity and is consistent with a PCoA analysis of the pilot study (Supplementary Figure S5), where the samples did not cluster by variety.

### Detection of multiple lineages of ***E. turcicum*** in infected leaves

We identified the genetic lineages of *E. turcicum* in the leaf samples by mapping all reads that Kaiju assigned to the genus *Exserohilum* on the *E. turcicum* reference genome. Out of 241 metagenomic sequencing samples, 41 had sufficient sequencing coverage of the *E. turcicum* genome (Materials and Methods) after mapping, filtering, and SNP calling. This set includes 26 samples from 2016, 4 samples from 2017, which correspond to the 30 samples with ≥ 70% of genome coverage in the direct mapping analysis (Fig. 2), all 11 samples from the pilot study (nine metagenomic samples and two isolates as controls), and no samples from 2018. This metagenomic SNP dataset was merged with 121 sequenced isolates from Europe and Kenya from ***Vidal-Villarejo et al.*** (***2023***) resulting in a final SNP dataset that included 10,098 SNPs and 162 samples.

Prior to analyzing the geographical distribution of pathogen diversity, we conducted tests to investigate the presence of multiple infections by different strains within individual samples. Initially, we employed a method that involved mapping reads to *MAT1-1* and *MAT1-2* sequences to identify the presence of both mating types within each sample. This analysis revealed the presence of *MAT1-1* in 33 samples, *MAT1-2* in 5 samples, and three samples with both mating types (Fig. 6A). Subsequently, we employed a second approach to assess the level of heterozygosity within *E. turcicum* samples obtained from leaves. Three samples exhibited nearly ten times the percentage of heterozygous sites compared to the two isolates used as control and they also harbored both *MAT1-1* and *MAT1-2* sequences (Fig. 6A). The pronounced heterozygosity observed in these samples, combined with the presence of both mating types, strongly indicates the occurrence of multiple infections by lineages belonging to different mating types.

**Figure 6.**
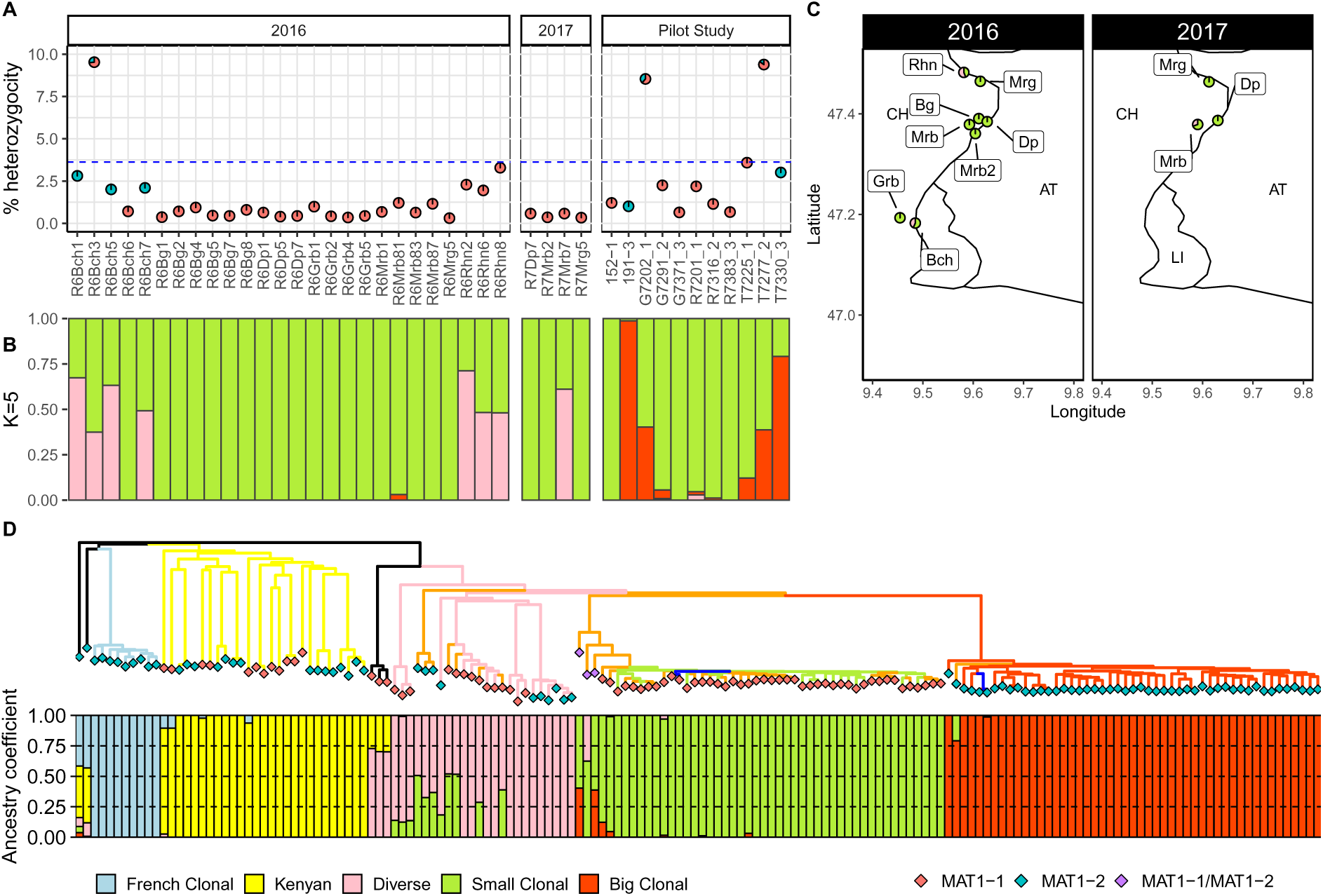
A) Percentage of heterozygosity shown as the percentage of heterozygous sites in the sample, and percentage of reads mapping to *MAT1-1* and *MAT1-2* sequences of the mating type locus, represented as pie charts. The dashed blue line in A indicates three times the percentage of the 152-1 isolate. B) Ancestry coefficients of *E. exserohilum* genomes in the metagenomic samples for 𝐾 = 5 clusters of the metagenomic samples and the two control isolates of the pilot study (152-1 and 191-3), with samples in the same order as in A. C) Distribution of ancestry coefficients in the sampling locations (agricultural fields). Pie charts summarize ancestry coefficients for all samples from one location. D) Ancestry coefficients for 𝐾 = 5 and the Neighbour-Joining tree for both metagenomic samples and isolates from Europe and Kenya ***Vidal-Villarejo et al.*** (***2023***). The metagenomic samples are indicated on the tree by orange branch colour and the two controls in the pilot study are indicated by dark blue branch colour. The branches of the isolates from Europe and Kenya are colored according to the genetic clusters as in ***Vidal-Villarejo et al.*** (***2023***). Rhombuses indicate the mating type, and the purple colour indicates that both mating types are present.

### Diversity and geographic distribution of ***E. turcicum*** lineages in the Swiss Rhine Valley

To assign the 41 samples with sufficient sequencing reads from *E. turcicum* to known genetic lineages, we utilized ADMIXTURE (***Alexander and Lange, 2011***) to estimate ancestral population coefficients. Additionally, we constructed a Neighbour-Joining (NJ) tree that incorporated previously sequenced isolates from Europe and Kenya, which served as a reference for characterized lineages in Europe. Out of the 41 metagenomic samples, the majority (28 samples; 68%) belonged to the Small Clonal lineage, as defined by ***Vidal-Villarejo et al.*** (***2023***). These samples displayed a high ancestry coefficient (>95%) of the Small Clonal cluster, clustered together with isolates from that lineage in the NJ Tree (Figs. 6B-D), and exclusively contained *MAT1-1* sequences. It is important to note that one of the 28 samples corresponds to the control sample (isolate 152-1) from the pilot study, which is a confirmed sample of the Small Clonal cluster.

A small number of six samples from 2016 and one from 2017 have only one mating type and were assigned by ADMIXTURE (with 𝐾 = 5) to the Diverse and Small Clonal lineages. Four of these samples group with the Diverse cluster in the NJ tree and their mating type is consistent with the mating type of the isolates in this sub-cluster (*MAT1-1*). The other three samples with the *MAT1-2* mating type form a separate group within the Diverse cluster, consistent with the overall structure of this cluster, because it consists of recently originated clonal lineages of the same mating type (***Vidal-Villarejo et al., 2023***). Within the pilot study, some samples also show Big Clonal ancestry coefficients, one of them belonging to the other isolate control, 191-3, which is a confirmed sample of the Big Clonal cluster.

The three samples (one from 2016 and two from the pilot study) with a very high number of heterozygous sites and both *MAT1-1* and *MAT1-2* alleles appear with admixed ancestries in ADMIXTURE, which might reflect either the presence of diploid individuals resulting from sexual recombination between different lineages, or a co-infection, where two distinct pathogenic lineages, each of a different mating type, inhabit the same host simultaneously. Given that European clonal clusters are associated with either one of the two mating types, we can directly estimate the proportion of infection for each lineage to determine if multiple infections occur in equal proportions or if one lineage dominates over the others. Specifically, for the three samples with evidence of multiple infections (R6Bch3, G7202_1, and T7277_2), the proportion of reads mapping to *MAT1-1* and *MAT1-2* mating types was 73.4% and 26.6% for R6Bch3, 61.3% and 38.7% for G7202_1, and 84.6% and 15.4% for T7277_2, respectively. Therefore, G7202_1 and T7277_2 consist of 61.3% and 84.6% Small Clonal (*MAT1-1*) infections, respectively, and of 38.7% and 15.4% Big Clonal (*MAT1-2*) infections, respectively. The ADMIXTURE analysis for sample R6Bch3 suggests an infection with the Small Clonal lineage and the Diverse cluster. The latter contains lineages of both mating types. Given that admixture coefficients for R6Bch3 are lower for Diverse than for Small Clonal lineages (Fig. 6B), we can infer that the proportion of *MAT1-1* sequencing reads (73.4%) in this sample belongs to the Small Clonal lineage rather than the Diverse lineage, which corresponds to *MAT1-2* (26.6%). Taken together, these three samples suggest that either co-infection rate or post-infection growth of the Small Clonal lineage is higher than for the other lineages.

### Local versus regional sources of ***E. turcicum*** infection

To determine whether local inoculum of *E. turcicum* infections originates from soil-borne spores or is distributed with annually and centrally produced seed stock, we analyzed samples from the most abundant Small Clonal lineage to recognize spatial differentiation of this lineage within the Rhine Valley region. This analysis included only the Small Clonal cluster because of a low sample size of the other clusters. The Small Clonal samples and one isolate from Kenya used as outgroup included 2,439 polymorphic SNPs, of which 872 segregate within Small Clonal samples. A NJ tree that was rooted with the sample from Kenya shows that samples from geographically closer sampling sites tend to cluster together and therefore are genetically more similar. (Fig. 7). A clear example are samples from Grabs (Grb), which cluster together with high confidence. Another example are some of the samples collected in 2016 from Balgach (Bg), Diepoldsau (Dp) and Marbach (Mrb) that cluster together, suggesting that genotypes from this lineage sampled in nearby locations are genetically more similar, although the bootstrap support is low. A comparison between the isolates and metagenomic samples belonging to the Small Clonal cluster did not exhibit a strong differentiation (e.g., by location and sampling year), which suggests that rapid divergence within clonal lineages is not ongoing. Overall, the analysis indicates a local clustering of lineages is more consistent with local soil-borne inoculation with *E. turcicum*, rather than seed-borne distribution of the pathogen by the centralised seed production this landrace.

**Figure 7.**
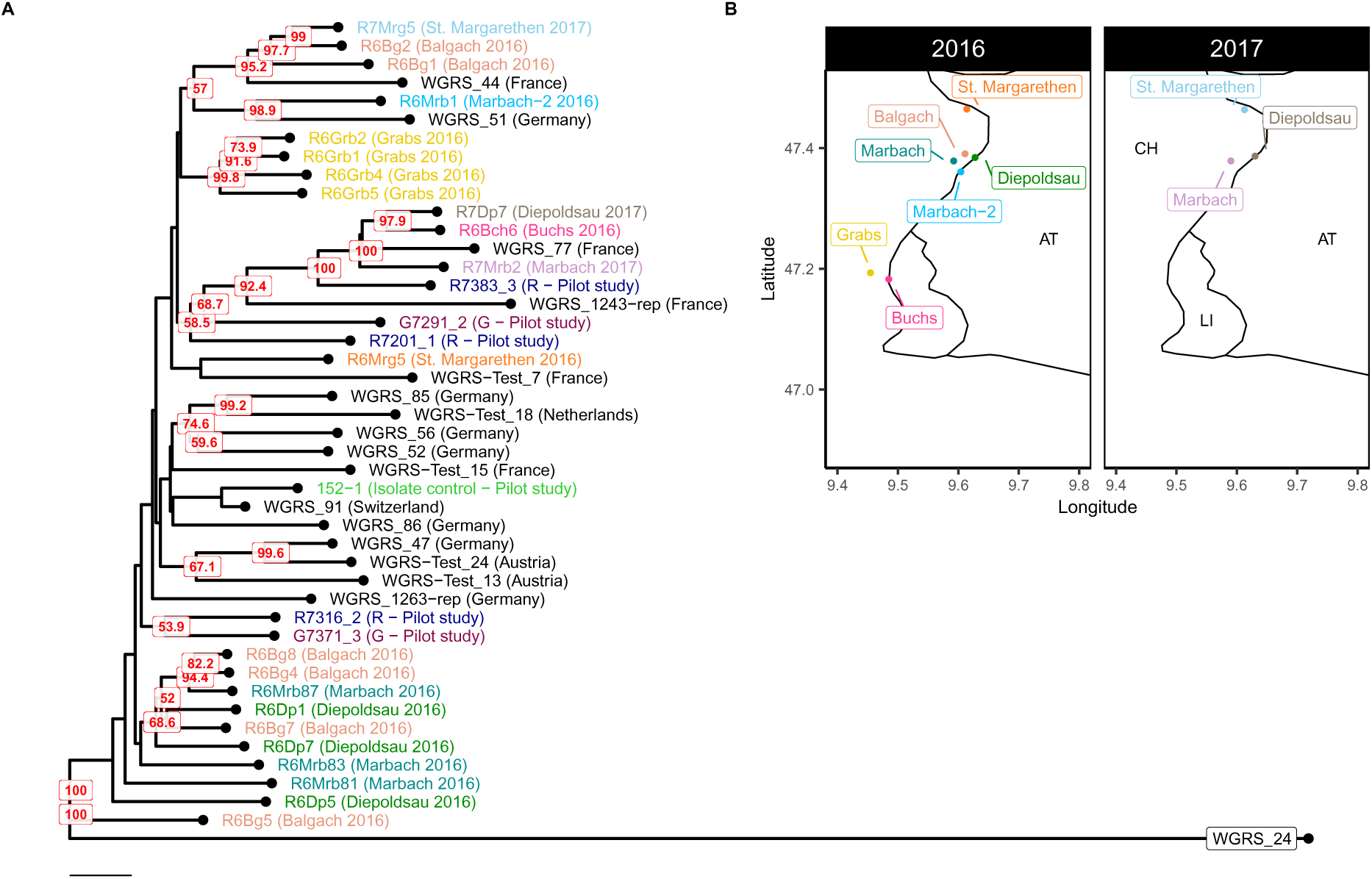
A) Neighbour-Joining tree of the Small Clonal samples rooted using one sample from Kenya (WGRS_24). Bootstrap values > 50% over 1,000 replicates are shown in red. Names are colored according to collection site (or the hybrid maize variety in the pilot study). European isolates from ***Vidal-Villarejo et al.*** (***2023***) are shown in black letters. The scale represents 1 unit of Euclidean distance. B) Geographic location of collection sites in 2016 and 2017.

### Co-occurrence network analysis of the microbial phyllobiome

The variation in microbial diversity prompted us to investigate correlations between the occurrence of taxa that may reflect antagonistic or synergistic interactions within the phyllobiome. Since correlation analyses, such as Pearson correlation, on compositional data (i.e., relative abundances) can be misleading due to the interdependence of percentages, we applied data transformations designed for compositional data analysis to overcome the challenges associated with interpreting raw relative abundances. To identify interactions between genera within the phyllobiome at the genus level, which may indicate communities with varying interaction strengths, we employed SPIEC-EASI (***Kurtz et al., 2015***). However, since the edge weights derived from SPIEC-EASI are not directly comparable to correlation coefficients, we utilized SparCC (***Friedman and Alm, 2012***) to weigh the relationships detected by SPIEC-EASI.

For a global interpretation we reduced the analysis to the top 100 most abundant genera to simplify the network. In addition, we analysed the complete dataset to focus on the interactions of all genera with *Exserohilum*. We produced two networks using two different methods implemented in SPIEC-EASI: neighborhood selection (MB method) and inverse covariance selection (glasso method). Both networks (Fig. 8A and B) show a clear division between Bacteria and Eukarya. The bacterial genera form two distinct clusters, within which the correlations are strongly positive. Between the two clusters, the connections are negative. The Eukarya also form two distinct clusters with strongly positive correlations among taxa within each cluster, but negative correlations between both clusters. Overall, four main groups with correlated abundances on the genus level can be identified.

**Figure 8.**
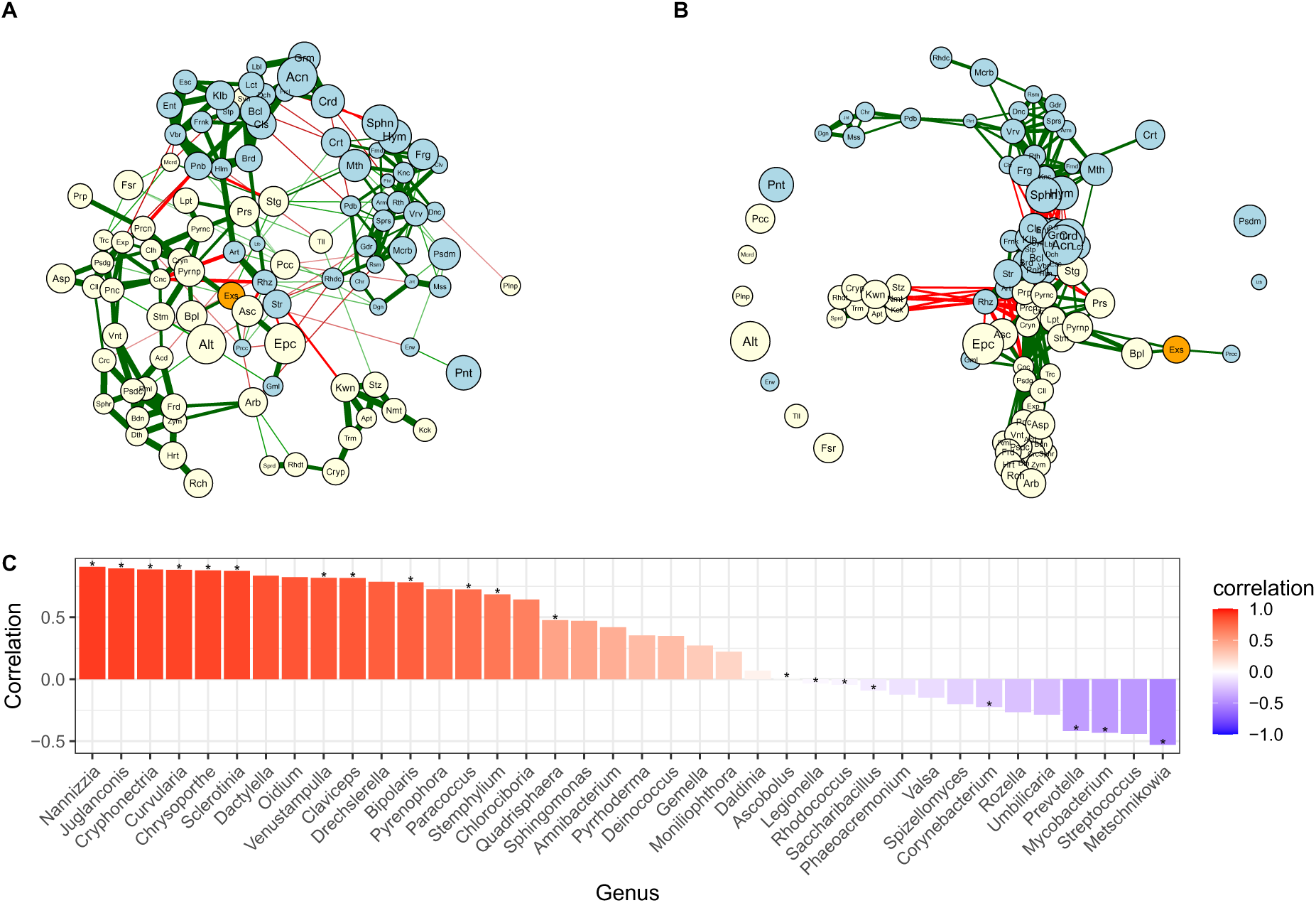
A) Co-occurrence network of the top 100 most abundant genera, generated using SPIEC-EASI with neighborhood selection and B) Inverse covariance selection, both weighted with SparCC. Genera are color-coded, with *Exserohilum* highlighted in orange, Eukarya in yellow, and Bacteria in blue. Negative correlations are depicted in red, while positive correlations are shown in green. The width of the relationship lines increases with the absolute correlation value. To enhance readability, the genera names have been shortened to three letters. C) SparCC correlation coefficients with *Exserohilum*. The asterisk (*) denotes the interaction present in both the Neighborhood and Inverse Covariance methods of SPIEC-EASI.

The position of *E. turcicum* in the networks created by the two methods indicates that this pathogen does not function as a strongly connected hub but rather occupies a marginal position (Fig. 8A and B). Among the 100 most frequent taxa, *Exserohilum* shows direct positive connections with *Bipolaris* (Ascomycota) and *Paracoccus* (Proteobacteria). Furthermore, when analyzing the complete dataset, *Exserohilum* exhibits positive correlations with other low-abundance genera (Fig. 8C). The strongest positive correlation observed across all samples is with *Nannizzia* (SparCC correlation coefficient, 𝑟 = 0.91), while the most negative correlation is with the genus *Metschnikowia* (𝑟 = −0.53). The negative direct interaction between *Exserohilum* and *Metschnikowia* is specifically observed in 2017 when analyzing the dataset for each year separately (Supplementary Fig. S6). However, in 2016 and 2018, there is a negative correlation according to SparCC, but this correlation is not identified as a direct interaction between *Exserohilum* and *Metschnikowia* in SPIEC-EASI (Supplementary Table 2). This suggests a possible indirect and negative interaction between the two genera.

### Influence of *E. turcicum* lineages on phyllobiome composition

To investigate the potential impact of different lineages of *E. turcicum* on the phyllobiome composition of individual leaves, we performed subsampling of the phyllobiome composition using 30 samples that exhibited sufficient coverage of *E. turcicum*. Subsequently, we plotted the dissimilarities in phyllobiome composition against the admixture ancestry coefficients (Fig. 9). The PCoA analysis of this subset revealed that samples assigned to the Diverse cluster were positioned on the left side of the plot, while samples with more than 90% Small Clonal cluster ancestry coefficients were dispersed throughout the plot. To determine whether the observed distribution of the Diverse cluster in the PCoA analysis could result from the influence of *E. turcicum* lineages, field location, or the proportion of ancestral coefficient of the Diverse cluster, we conducted a PERMANOVA on the subset. We used the collection site and the ancestry coefficients of the Diverse cluster as factors and stratified by year. The analysis indicated that the collection site does not influence significatively the variation in phyllobiome composition (𝑅^2^ = 0.345, 𝑝 = 0.05) nor does the ancestry coefficient of the Diverse cluster (𝑅^2^ = 0.031, 𝑝 = 0.33) in this subset.

**Figure 9.**
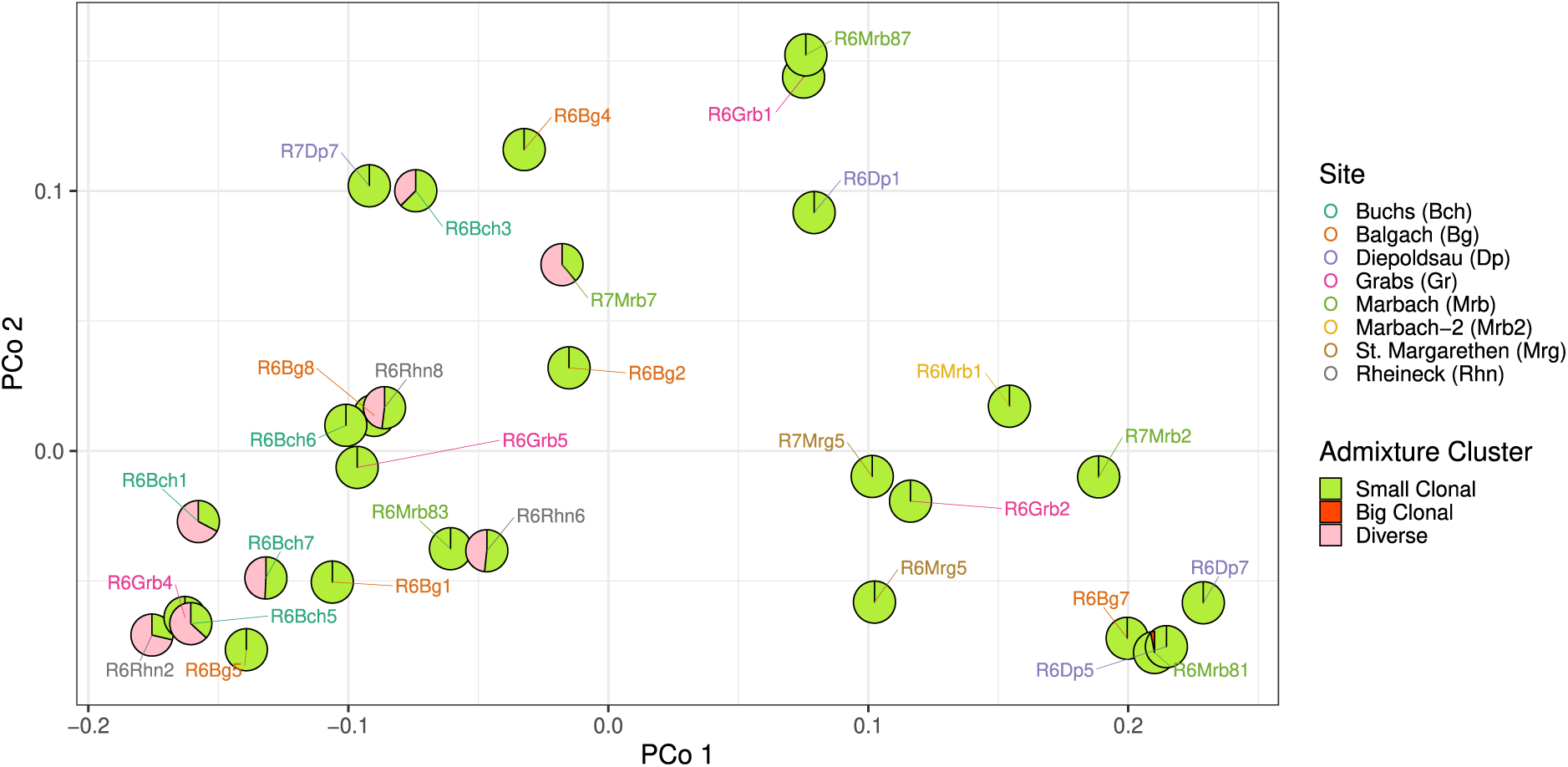
PCoA plot showing the dissimilarities in phyllobiome composition among samples with sufficient *E. turcicum* coverage, along with their corresponding ADMIXTURE ancestry coefficients (𝐾 = 5). The samples are labeled with their names and colored according to their collection site.

## Discussion

Our metagenomic survey of maize leaves infected with the fungal pathogen *E. turcicum* revealed highly variable numbers of sequencing reads from this pathogen between samples. The relative proportion of maize DNA was significantly higher than that of *E. turcicum* DNA among the sequenced samples, exhibiting substantial variation across years, locations, and individual samples. Out of 241 samples, only 41 (17%) contained a sufficient number of *E. turcicum* reads for subsequent analyses. The Kaiju classifier successfully classified 32% of all sequenced reads at the genus level, whereas 57% of the reads were unclassified (which include all maize reads), and 10% could not be assigned to a specific genus. We conclude that the identification and sampling of infected leaves needs to be optimized for a robust large-scale metagenomic monitoring of pathogen genetic diversity.

Despite this limitation, we observed substantial genetic diversity both within and between farmer’s fields in the Swiss Rhine valley. The majority of the samples (38 out of 41) represent single infections of the pathogen, which was expected given that we aimed at extracting DNA from a single lesion per leaf only. One advantage of metagenomic sequencing of single infections is that it allows for more reliable determination of the genetic lineage of *E. turcicum* and enables investigation of its interaction with the accompanying phyllobiome of the infected leaf area with higher statistical quality.

Based on our previous study of European isolates (***Vidal-Villarejo et al., 2023***) we were able to assign all metagenomic samples to known *E. turcicum* clonal lineages. Given the clonality of most lineages and their different mating types, independent infections in leaves with multiple infection could be quantified by using the percentage of reads mapping to each mating type. The most common lineage among samples from the Rhine Valley is the so called Small Clonal lineage which is represented by the *MAT1-1* mating type. We did not observe new lineages in our sample, indicating that since the collection of the isolates in 2011 and 2012, no new lineages seem to have arrived or arisen (e.g., by sexual recombination) in the Swiss Rhine valley. Three samples showed clear evidence of multiple infections of the same maize leave with spores from different lineages and/or an infection of recent recombinants between different lineages. Since recombination of *E. turcicum* genotypes has not been previously observed in Europe and is unlikely to occur in temperate regions (***Borchardt et al., 1998***; ***Vidal-Villarejo et al., 2023***), an infection by recent recombinants seems unlikely and the presence of different mating types in a single leaf sample likely corresponds to multiple infections of different clonal lineages. However, if suitable conditions for *E. turcicum* to reproduce sexually were to be met in Europe (e.g., high temperatures and humidity due to global climate change), sexual recombination may lead to an increase in genetic variation and enhance virulence or the appearance of new resistant varieties (***Bunkoed et al., 2014***).

The phylogenetic tree of Small Clonal samples reflects geographic clustering, where samples from the same or nearby location are clustered together. This suggests that the inoculum originates mainly from local soild-borne spores rather than from the annual, centralized distribution of seed stocks.

### High variation in diversity of the maize phyllobiome

Sequencing of phyllobiome revealed that both sampling year and field location, as well as their interaction, are associated with the dissimilarities observed of microbial composition, while North-South direction of the Rhine Valley has no significant influence. Notably, the phyllobiome composition in 2018 exhibited substantial differences compared to the previous two years, likely due to distinct weather conditions characterized by lower precipitation and higher mean temperatures. The phyllobiome, being directly exposed to the environment, is particularly susceptible to the impact of rapidly fluctuating conditions, which may have a more pronounced effect on leaf surface communities compared to those inhabiting internal plant compartments (***Trivedi et al., 2020***, ***2022***). For example, ***Aydogan et al. (2018)*** reported a reduced abundance of *Sphingomonas spp*. and *Rhizobium spp*. in the phyllobiome of the annual herb *Galium album* in experimental plots with a higher temperature, along with an increased abundance of Enterobacteriaceae (including *Acinetobacter*). These findings are consistent with our study, because we observed a higher relative abundance of *Acinetobacter* in 2018 compared to the two previous, colder years, along with a lower presence of *Sphingomonas*. However, the variation in phyllobiome composition across all field locations cannot be solely attributed to year effects. Specifically, we found no significant effect of year on phyllobiome composition in St. Margarethen and Grabs, as indicated by the non-significant results of the PERMANOVA tests. This suggests that factors other than year, such as temperature, sampling date, plant developmental stage, fertilizers, and soil characteristics, may be driving the observed differences in phyllobiome composition (***Van Overbeek and Van Elsas, 2008***; ***Bulgari et al., 2014***; ***Bokulich et al., 2014***; ***Coince et al., 2014***; ***Robinson et al., 2016***).

Among the most abundant genera identified in the phyllobiome, several were common plant endophytes such as *Alternaria*, *Epicoccum*, *Acinetobacter*, *Pseudomonas*, *Fusarium*, *Pantoea*, and *Sphingomonas* (***Fisher et al., 1992***; ***Liu et al., 2012***; ***Kandel et al., 2017***; ***CAB International, 2020***). In a few samples, the dominant genus was *Puccinia*, the causal agent of common rust of maize (***CAB International, 2020***). Additionally, *Bipolaris*, the causal agent of Southern Corn Leaf Blight (***CAB International, 2020***), exhibited high abundance in only two samples. No other putative pathogens were found in high abundance, except for *Exserohilum*, which was present in some samples but not as the most dominant genus overall. Despite sampling leaves from visibly infected plants with *E. turcicum*, the genus *Exserohilum* was not the most dominant in all samples. Several explanations can be suggested for the observed low abundance of *Exserohilum*. First, the lower humidity in 2018 could account for the reduced presence of *E. turcicum* during that year, as this pathogen is known to prefer high humidity. Second, misidentification of plants displaying Northern Corn Leaf Blight (NCLB) symptoms could have occurred, particularly in samples where *Puccinia* or *Bipolaris* were dominant, but not in others where no other pathogen seemed to be prevalent. Third, delayed sample processing after infection may have allowed saprophytic organisms to become active, potentially masking the primary infection as the leaf tissue deteriorates or becomes necrotic. This could explain the samples with high abundance of *Epicoccum* (***Schol-Schwarz, 1959***; ***Abdel-Azeem et al., 2021***). However, to prevent such a scenario, we immediately after harvest dried leaf samples with silica gel and stored them at cool conditions.

### Identification of potential biological control agents

We used co-occurrence network analysis in all 241 samples to investigate possible interactions between *E. turcicum* and other taxa of the phyllobiome, and we identified four groups of genera that are highly correlated (either negatively or positively). Among these groups, the one containing the *Acinetobacter* genus showed the most negative correlations with the other groups, specially with the bacterial group containing *Sphingomonas*. Furthermore, our analysis revealed the potential of *Metschnikowia* as Biological Control Agent (BCA). This potential arises from its strong negative correlation with *Exserohilum*, its non-pathogenic nature towards maize and other plants, and from previous research that has investigated its suitability as a BCA (***Manso and Nunes, 2011***). Additionally, we identified *Corynebacterium* as another negatively correlated genus with *Exserohilum*, which has been previously tested as a BCA for this pathogen (***Sartori et al., 2015***, ***2017***). In summary, these results demonstrate the efficacy of network analysis in identifying potential BCAs for *Exserohilum*.

### Improving the design of monitoring studies

Although the above results show the value of pathogen monitoring by sequencing, our study has some limitations in its design that should be addressed in future efforts. First, the highly variable proportion of *E. turcicum* DNA in individual leaf samples indicates that one needs to reliably identify leaves with *E. turcicum* infections and, if possible, sample only single infections in the leaf. Second, sampling over multiple years allows an analysis of environmental effects on pathogen diversity and pathogen-leaf microbiome interactions, while appropriate controls such as sequencing uninfected plants for the microbiome should be used. Third, comparisons among different maize varieties can help to explore the effect of the host genetic background in natural infections, but such effects may be difficult to evaluate with small sample sizes or an appropriate design of field trials. Fourth, pathogen monitoring may benefit from on site long read sequencing with portable sequencing devices in combination with experimental or computational pathogen enrichment techniques (***Johnson et al., 2023***). Overall, we demonstrated that metagenome analysis of infected leaves is a suitable alternative to the extraction and sequencing of single isolates for pathogen monitoring for the management of pathogen resistances and improving resistance breeding.

## Material and Methods

### Leaf sampling and metagenome sequencing

A team of workers conducted sampling of all fields every year within three days. The sampling strategy entailed moving through the field in a transect and collecting a single leaf sample after visual inspection for an *E. turcicum* infection. Approximately 10 leaves were sampled per field. Samples were assigned to the same collection site (agricultural farm, denoted with the name of the town) when they came from the same town and the distance of the samples within site was smaller than with samples of other sites. Whenever a sample had a higher distance with samples from the same town than samples from nearby towns, it was assigned to a different collection site (denoted with the suffix ‘2‘ in the name of the site). The sampled leaves were cut out with scissors and stored in teabags, which were then placed in closable plastic bags with dried silica gel beads to quickly desiccate the leaves and prevent post-sampling degradation. The dried leaves were cut into smaller pieces with scissors and placed in the wells of a deep well 96 plate. Six ceramic beads (2.8 mm diameter; MoBio, USA) were added to each well, and the tissue was ground in a Retsch mixer mill (MM400) for 30 seconds at a speed of 30 sec^−1^. The DNA was extracted using the Micro AX Blood Gravity kit (A&A Biotechnology, Poland; Cat. No. 101-100) following the manufacturer’s protocol. The DNA was then diluted to a concentration of 2.5 ng µl^−1^ in EB buffer, and its quality was analyzed on an agarose gel. Whole genome sequencing libraries were created using a multiplex tagmentation protocol (***Baym et al., 2015***). The libraries were paired-end sequenced (2 x 150 bp) using a HiScan Illumina sequencer (Macrogen, South Korea).

### Metagenomic sequence processing and taxonomic classification

To process the raw sequencing reads and map them to the genomes of *E. turcicum* and maize, we developed a Snakemake pipeline The pipeline involved several steps, starting with read cleaning using TrimGalore (***Krueger, 2015***). The TrimGalore parameters used for this process were ‘–paired –fastqc –clip_R1 5 –clip_R2 5 –three_prime_clip_R1 5 –three_prime_clip_R2 5’. Following read cleaning, the processed reads were mapped to the *E. turcicum* genome using the bwa mem algorithm (***Li and Durbin, 2009***). To extract unmapped reads from the resulting BAM files, we employed samtools view (***Li et al., 2009***; ***Li, 2011***) with the ‘-f 4’ flag. Subsequently, the unmapped reads were converted back to FastQ format using picard samToFastq from the Picard tool suite (http://broadinstitute.github.io/picard/). Next, the unmapped reads were aligned to the maize B73 reference genome V4 (AGPv4) using the bwa mem algorithm. To assess mapping percentages for both *Exserohilum* and maize, we calculated mapping summaries utilizing Qualimap (***Okonechnikov et al., 2016***) as an integral part of the Snakemake pipeline. It is important to note that the sole purpose of this pipeline was to perform read cleaning and generate summary statistics pertaining to mapping percentages for *Exserohilum* and maize.

Following the initial mapping approach, we developed another Snakemake pipeline to taxonomically classify the previously cleaned reads using Kaiju (***Menzel et al., 2016***). To facilitate this classification, the pipeline generated the Kaiju database “nr_euk,” which is a subset of the NCBI BLAST nr protein database encompassing data from archaea, bacteria, viruses, fungi, and microbial eukaryotes. To obtain a genus-level summary of the Kaiju output, we employed the kaiju2table script integrated within the Kaiju program. Subsequently, we excluded six samples from our analysis due to a low number of sequencing reads, specifically those with fewer than 1,000 reads. Furthermore, we implemented a filtering step to eliminate false positives by removing eukaryotic and bacterial genera with a minimum relative abundance of less than 0.01% across fewer than 10 samples. This stringent filtering was necessary to mitigate the risk of an increased number of false positives, potentially ranging from tens to thousands, which has been reported in previous studies (***Ye et al., 2019***).

### Extraction of *E. turcicum* reads and variant calling

To obtain a representative sample of reads specific to *E. turcicum*, the Kaiju Snakemake pipeline implemented sub-sampling of all reads classified by Kaiju as belonging to taxonomic IDs 671987 (*Exserohilum turcicum* isolate Et28A) and 93612 (*Exserohilum turcicum*). This sub-sampling process utilized the seqtk program (***Li, 2013***). Subsequently, the resulting subset of reads was aligned to the reference genome of *Setosphaeria turcica* Et28A v1.0 (***Ohm et al., 2012***; ***Condon et al., 2013***) using the bwa-mem2 algorithm (***Vasimuddin et al., 2019***). To enhance the accuracy of the mapping process, we performed local realignment around InDels and eliminated duplicate reads using GATK (***McKenna et al., 2010***). To avoid incorrect assignment of reads originating from closely related species, we implemented a filtering step that discarded reads mapping with more than two mismatches. This filtering was carried out using the BamTools filter -tag ‘NM:<=2’ command (***Barnett et al., 2011***).

For Single Nucleotide Polymorphism (SNP) calling, we utilized SAMtools mpileup in combination with Bcftools call (***Narasimhan et al., 2016***). During this step, we implemented two filtering thresholds: a minimum read quality filter of 28 and a minimum mapping quality filter of 20. To ensure the reliability of the called SNPs as true positive *E. turcicum* SNPs, we specifically retained those SNPs that had been previously observed in our comprehensive survey of 121 individuals from Europe and Kenya (***Vidal-Villarejo et al., 2023***). Subsequently, we performed additional filtering by excluding samples with more than 50% of missing data and loci (genomic positions) with more than 40% of missing data, and we kept only bi-allelic SNPs.

### Identification of *E. turcicum* lineages in sequencing data

To assign the metagenomic *E. turcicum* sequencing reads to known genetic lineages of *E. turcicum* sequenced from infected leaf samples, we conducted several analyses. First, we mapped the *E. turcicum* reads to the two *MAT1* mating type locus sequences (GenBank accessions GU997138.1 and GU997137.1) and classified samples accordingly. We considered a minimum of 50% of reads mapping to the inner region of the *MAT1* sequences, which was taken from 500 bp to 1,700 bp in both sequences in order to consider a positive *MAT1-1/MAT1-2* type. Second, we used ADMIXTURE (***Alexander and Lange, 2011***) to obtain the ancestral groups of the *E. turcicum* dataset, including the set of 121 sequenced isolates (***Vidal-Villarejo et al., 2023***). We performed 20 independent runs with *K* ranging from 1 to 15, with the calculation of cross-validation error option activated (Supplementary Fig. S7) We then used CLUMPAK (***Kopelman et al., 2015***) to merge the independent runs into a single result, and we used MajorCluster (i.e. the cluster result containing most replicate runs, according to the authors of CLUMPAK) to represent the results. Third, we built a Neighbour-Joining tree with the Euclidean distances, of allele frequencies calculated from read counts, using R package ape (***Saitou and Nei, 1987***). Positions with < 2 reads were set as missing and an additional filtering of 20% of missing data per SNP was applied for the calculus of allele frequencies. In addition, we assessed the percentage of heterozygous genotypes in each sample using the same filtered set used in the NJ tree.

We then selected a subset of samples classified as Small Clonal lineage in ***Vidal-Villarejo et al.*** (***2023***) and constructed a new NJ tree based on this subset. To ensure the clonal nature of the samples, we only included those with a low proportion of heterozygous genotype calls (<2.5%), a single mating type, and little or no ancestry admixture with other clusters than Small Clonal. This analysis also incorporated the sequencing data of Small Clonal isolates from Europe and one sample from Kenya as outgroup in order to root the tree. Bootstrap values were calculated from 1,000 replicates with boot.phylo function from R package ape.

### Taxonomic diversity of infected leaf phyllobiome

We obtained the taxonomic diversity of the infected leaf phyllobiomes at the genus level using the kaiju2table summary. This summary table contains taxonomic read counts, which indicates the number of reads classified for each taxonomic ID encountered by Kaiju. To study the correlation between phyllobiome composition and time (years), site (collection sites) and latitudinal and longitudinal gradients (latitude and longitude), we performed a permutational analysis of variance (PERMANOVA) over the fourth variables using Bray-Curtis distances on the root squared transformation of the relative abundances (𝑓 𝑜𝑟𝑚𝑢𝑙𝑎 = 𝑏𝑐𝑢𝑟.𝑑𝑖𝑠𝑡 ∼ 𝑦𝑒𝑎𝑟 ∗ 𝑠𝑖𝑡𝑒+𝑙𝑎𝑡𝑖𝑡𝑢𝑑𝑒+𝑙𝑜𝑛𝑔𝑖𝑡𝑢𝑑𝑒) and 99,999 permutations. We carried out the PERMANOVA analysis with three different datasets: eukaryotic genera only, bacterial genera only, and both eukaryotic and bacterial genera combined. Therefore, the relative abundances were relative to the total number of reads classified as the superkingdom(s) being analyzed, not from the total number of raw reads. To ensure that significant results were not an artifact of different dispersions between the data, we performed a permutation test of homogeneity of the dispersion. When the homogeneity test showed significant dispersion, we subsampled the data with 20 random samples, 500 times, of each group and recalculated the PERMANOVA and the test of homogeneity of the dispersion. We calculated pairwise PERMANOVA by taking only the year as the factor (𝑓 𝑜𝑟𝑚𝑢𝑙𝑎 = 𝑏𝑐𝑢𝑟.𝑑𝑖𝑠𝑡 ∼ 𝑦𝑒𝑎𝑟). The PERMANOVA reduced to the samples with Small Clonal *E. turcicum* lineage was calculated with collection site and ancestry coefficients of Diverse cluster, stratifying by the year (𝑓 𝑜𝑟𝑚𝑢𝑙𝑎 = 𝑏𝑐𝑢𝑟.𝑑𝑖𝑠𝑡 ∼ 𝑠𝑖𝑡𝑒 + 𝑑𝑖𝑣𝑒𝑟𝑠𝑒_𝑎𝑛𝑐𝑒𝑠𝑡𝑟𝑦, 𝑠𝑡𝑟𝑎𝑡𝑎 = 𝑦𝑒𝑎𝑟). We performed Principal Coordinate Analysis (PCoA) to visualize the phyllobiome dissimilarities. To measure the eukaryotic and bacterial diversity of the samples, we calculated Shannon’s index, 𝐻, from the relative abundances. For all the aforementioned tests, we used the functions provided in the R package vegan (***Oksanen et al., 2019***). Paired t-test and Wilcoxon test in Shannon’s index 𝐻 were computed with the R package ggsignif (***Constantin and Patil, 2021***).

### Agrometereological data

Agrometeorological historical data was extracted from the portal http://www.agrometeo.ch (Accessed October 2020). We downloaded data by month from 2016 to 2018 including months from April to September and all meteorological stations in the Rheintal region, which includes: Bad Ragaz, Berneck-Feuerbrand, Berneck-Indermaur, Cazis, Fläsch, Frümsen, Grabs, Jenins, Kriessern, Landquart, Maienfeld, Malans-Obstbau, Malans-Weinbau, Salez, Sargans, Thal, Walenstadt, Weite and Zizers. To test for significant differences between the three years, we used the Wilcoxon test implemented in the R package ggsignif (***Constantin and Patil, 2021***).

### Microbial interaction networks

We calculated correlation networks of the observed genera using the R package SPIEC-EASI (***Kurtz et al., 2015***), which also implements a function to calculate SparCC correlation (***Friedman and Alm, 2012***). SPIEC-EASI utilizes a graphical model inference framework to reconstruct relationships among genera. Unlike other methods, SPIEC-EASI avoids detecting relationships between taxa that are correlated but indirectly connected (***Kurtz et al., 2015***), whereas SparCC calculates correlations which are directly comparable to correlation coefficients. Therefore we used SPIEC-EASI to identify truly direct relationships between taxa and SparCC to weight and interprete those relationships. Both SPIEC-EASI and SparCC apply data transformation designed for compositional data and assume that the true community network is sparse.

Both available SPIEC-EASI methods, mb (neighborhood selection) and glasso (inverse covariate), were calculated with 40 lambda and 50 repetitions, and a minimum of 0.01 for each method. The relationships were then weighted with correlation coefficients from SparCC. Raw read counts were used as input since the methods in this section perform their own data transformation. To simplify the interpretation of the networks with numerous nodes and edges, we calculated the interaction network with a subset of the 100 most abundant genera and displayed it with a spring layout using the R package qgraph (***Epskamp et al., 2012***). We also used a co-occurrence network calculated from the complete dataset (not only the 100 most abundant) to investigate the direct relationships and correlation coefficients with *Exserohilum* to account for all possible interacting genera with this pathogen.

## Author Contributions

M.V-V. and K.S designed the study. B.K, M.H., B.O and H.O designed leaf sampling strategy, collected and provided leaf material. B.D built the mapping Snakemake Pipeline. M.V-V built the Kaiju Snakemake Pipeline and conducted the data analyses. M.V-V and K.S. interpreted the data and wrote the manuscript. All authors read and agreed to the final version of the manuscript.

## Data Availability

Raw sequencing data were deposited to ENA under project number PRJEB66236. Derived sequencing data like vcf files were deposited and analysis scripts are available from Zenodo (https://10.5281/zenodo.10871178).

## Acknowledgments

We are grateful to Elisabeth Kokai-Kota for DNA extraction and sequencing library construction. This work was funded by a grant in the DFG Priority Program SPP1819 “Rapid evolutionary adaptation: Potential and constraints” (SCHM 1354-11/1) and supported by the Bundesministerium für Bildung und Forschung (BMBF) under grant number 031B0731A. The article processing charges (APC) were also covered by the Bundesministerium für Bildung und Forschung (BMBF) under grant number 031B0731A. Additional funding was provided from the Swiss National Action Plan NAP-PGREL project PGRL-NN-0061 to H.O. and B. O. The authors acknowledge the additional support by the state of Baden-Württemberg through bwHPC.

## Notes

### Competing Interest Statement

The authors have declared no competing interest.

## References

Abdel-Azeem AM, Yadav AN, Yadav N, Usmani Z. Industrially Important Fungi for Sustainable Development: Volume 1: Biodiversity and Ecological Perspectives. Springer; 2021.

Alexander DH, Lange K. Enhancements to the ADMIXTURE algorithm for individual ancestry estimation. BMC Bioinformatics. 2011; 12. doi: 10.1186/1471-2105-12-246.

Aydogan EL, Moser G, Müller C, Kämpfer P, Glaeser SP, Long-Term Warming Shifts the Composition of Bacterial Communities in the Phyllosphere of Galium album in a Permanent Grassland Field-Experiment; 2018. https://www.frontiersin.org/articles/10.3389/fmicb.2018.00144.

Barnett DW, Garrison EK, Quinlan AR, Strömberg MP, Marth GT. BamTools: a C++ API and toolkit for analyzing and managing BAM files. Bioinformatics. 2011 04; 27(12):1691–1692. 10.1093/bioinformatics/btr174, doi: 10.1093/bioinformatics/btr174.

Baym M, Kryazhimskiy S, Lieberman TD, Chung H, Desai MM, Kishony R. Inexpensive Multiplexed Library Preparation for Megabase-Sized Genomes. PLOS ONE. 2015 May; 10(5):e0128036. http://dx.plos.org/10.1371/journal.pone.0128036, doi: 10.1371/journal.pone.0128036.

Bokulich NA, Thorngate JH, Richardson PM, Mills DA. Microbial biogeography of wine grapes is conditioned by cultivar, vintage, and climate. Proceedings of the National Academy of Sciences of the United States of America. 2014; 111(1):139–148. doi: 10.1073/pnas.1317377110.

Borchardt DS, Welz HG, Geiger HH. Genetic structure of Setosphaeria turcica populations in tropical and temperate climates. Phytopathology. 1998; 88(4):322–329. doi: 10.1094/PHYTO.1998.88.4.322.

Bulgari D, Casati P, Quaglino F, Bianco PA. Endophytic bacterial community of grapevine leaves influenced by sampling date and phytoplasma infection process. BMC Microbiology. 2014; 14(1):1–11. doi: 10.1186/1471-2180-14-198.

Bunkoed W, Kasam S, Chaijuckam P, Yhamsoongnern J, Prathuangwong S. Sexual reproduction of setosphaeria turcica in natural corn fields in Thailand. Kasetsart Journal Natural Science. 2014; 48(2):175–182.

CAB International, Invasive Species Compendium; 2020. https://www.cabi.org/isc.

Chaloner TM, Gurr SJ, Bebber DP. Plant pathogen infection risk tracks global crop yields under climate change. Nature Climate Change. 2021 Aug; 11(8):710–715. https://www.nature.com/articles/s41558-021-01104-8, doi: 10.1038/s41558-021-01104-8, number: 8 Publisher: Nature Publishing Group.

Coince A, Cordier T, Lengellé J, Defossez E, Vacher C, Robin C, Buée M, Marçais B. Leaf and root-associated fungal assemblages do not follow similar elevational diversity patterns. PLoS ONE. 2014; 9(6). doi: 10.1371/journal.pone.0100668.

Condon BJ, Leng Y, Wu D, Bushley KE, Ohm RA, Otillar R, Martin J, Schackwitz W, Grimwood J, MohdZainudin N, Xue C, Wang R, Manning VA, Dhillon B, Tu ZJ, Steffenson BJ, Salamov A, Sun H, Lowry S, LaButti K, et al. Comparative Genome Structure, Secondary Metabolite, and Effector Coding Capacity across Cochliobolus Pathogens. PLOS Genetics. 2013 01; 9(1):1–29. 10.1371/journal.pgen.1003233, doi: 10.1371/journal.pgen.1003233.

Constantin AE, Patil I. ggsignif: R Package for Displaying Significance Brackets for ‘ggplot2’. PsyArxiv. 2021; https://psyarxiv.com/7awm6, doi: 10.31234/osf.io/7awm6.

Epskamp S, Cramer AOJ, Waldorp LJ, Schmittmann VD, Borsboom D. qgraph: Network Visualizations of Relationships in Psychometric Data. Journal of Statistical Software. 2012; 48(4):1–18. http://www.jstatsoft.org/v48/i04/.

Fisher PJ, Petrini O, Scott HML. The distribution of some fungal and bacterial endophytes in maize (Zea mays L.). New Phytologist. 1992; 122(2):299–305. doi: 10.1111/j.1469-8137.1992.tb04234.x.

Friedman J, Alm EJ. Inferring Correlation Networks from Genomic Survey Data. PLoS Computational Biology. 2012; 8(9):1–11. doi: 10.1371/journal.pcbi.1002687.

Galiano-Carneiro AL, Miedaner T. Genetics of Resistance and Pathogenicity in the Maize/Setosphaeria Turcica Pathosystem and Implications for Breeding. Frontiers in Plant Science. 2017; 8. https://www.frontiersin.org/articles/10.3389/fpls.2017.01490/full, doi: 10.3389/fpls.2017.01490.

Hanekamp H. Europäisches Rassen-Monitoring und Pathogenesestudien zur Turcicum-Blattdürre (*Exserohilum turcicum*) an Mais (*Zea mays* L.). Ph.D. Thesis, University of Göttingen; 2016.

Hubbard A, Lewis CM, Yoshida K, Ramirez-Gonzalez RH, de Vallavieille-Pope C, Thomas J, Kamoun S, Bayles R, Uauy C, Saunders DG. Field Pathogenomics Reveals the Emergence of a Diverse Wheat Yellow Rust Population. Genome Biology. 2015 Dec; 16(1). http://genomebiology.com/2015/16/1/23, doi: 10.1186/s13059-015-0590-8.

Johnson MA, Vinatzer BA, Li S. Reference-Free Plant Disease Detection Using Machine Learning and Long-Read Metagenomic Sequencing. Applied and Environmental Microbiology. 2023 May; 89(6):e00260–23. https://journals.asm.org/doi/full/10.1128/aem.00260-23, doi: 10.1128/aem.00260-23, publisher: American Society for Microbiology.

Kandel S, Joubert P, Doty S. Bacterial Endophyte Colonization and Distribution within Plants. Microorganisms. 2017; 5(4):77. doi: 10.3390/microorganisms5040077.

Karasov TL, Neumann M, Duque-Jaramillo A, Kersten S, Bezrukov I, Schröppel B, Symeonidi E, Lundberg DS, Regalado J, Shirsekar G, Bergelson J, Weigel D, The relationship between microbial biomass and disease in the Arabidopsis thaliana phyllosphere. bioRxiv; 2019. https://www. biorxiv.org/content/10.1101/828814v1, doi: 10.1101/828814, pages: 828814 Section: New Results.

Kopelman NM, Mayzel J, Jakobsson M, Rosenberg NA, Mayrose I. Clumpak: a program for identifying clustering modes and packaging population structure inferences across K. Molecular Ecology Resources. 2015; 15(5):1179–1191. https://onlinelibrary.wiley.com/doi/abs/10.1111/1755-0998.12387, doi: 10.1111/1755-0998.12387.

Krueger F, Trim Galore! A wrapper tool around Cutadapt and FastQC to consistently apply quality and adapter trimming to FastQ files; 2015. http://www.bioinformatics.babraham.ac.uk/projects/trim_galore/.

Kurtz ZD, Müller CL, Miraldi ER, Littman DR, Blaser MJ, Bonneau RA. Sparse and Compositionally Robust Inference of Microbial Ecological Networks. PLoS Computational Biology. 2015; 11(5):1–25. doi: 10.1371/journal.pcbi.1004226.

Li H. A statistical framework for SNP calling, mutation discovery, association mapping and population genetical parameter estimation from sequencing data. Bioinformatics. 2011; 27(21):2987– 2993. 10.1093/bioinformatics/btr509, doi: 10.1093/bioinformatics/btr509.

Li H, Seqtk: a fast and lightweight tool for processing FASTA or FASTQ sequences; 2013. https://github.com/lh3/seqtk.

Li H, Durbin R. Fast and accurate short read alignment with Burrows–Wheeler transform. Bioinformatics. 2009; 25(14):1754–1760. +10.1093/bioinformatics/btp324, doi: 10.1093/bioinformatics/btp324.

Li H, Handsaker B, Wysoker A, Fennell T, Ruan J, Homer N, Marth G, Abecasis G, Durbin R, Subgroup GPDP. The Sequence Alignment/Map format and SAMtools. Bioinformatics. 2009; 25(16):2078– 2079. 10.1093/bioinformatics/btp352, doi: 10.1093/bioinformatics/btp352.

Liu Y, Zuo S, Xu L, Zou Y, Song W. Study on diversity of endophytic bacterial communities in seeds of hybrid maize and their parental lines. Archives of Microbiology. 2012; 194(12):1001–1012. doi: 10.1007/s00203-012-0836-8.

Manso T, Nunes C. Metschnikowia andauensis as a new biocontrol agent of fruit postharvest diseases. Postharvest Biology and Technology. 2011; 61(1):64–71. 10.1016/j.postharvbio.2011.02.004, doi: 10.1016/j.postharvbio.2011.02.004.

McDonald BA, Stukenbrock EH. Rapid Emergence of Pathogens in Agro-Ecosystems: Global Threats to Agricultural Sustainability and Food Security. Philosophical Transactions of the Royal Society B: Biological Sciences. 2016 Dec; 371(1709):20160026. http://rstb.royalsocietypublishing.org/lookup/doi/10.1098/rstb.2016.0026, doi: 10.1098/rstb.2016.0026.

McKenna A, Hanna M, Banks E, Sivachenko A, Cibulskis K, Kernytsky A, Garimella K, Altshuler D, Gabriel S, Daly M, DePristo MA. The Genome Analysis Toolkit: A MapReduce framework for analyzing next-generation DNA sequencing data. Genome Research. 2010; 20(9):1297–1303. http://genome.cshlp.org/content/20/9/1297.abstract, doi: 10.1101/gr.107524.110.

Menzel P, Ng KL, Krogh A. Fast and sensitive taxonomic classification for metagenomics with Kaiju. Nature Communications. 2016; 7:11257. http://www.nature.com/doifinder/10.1038/ncomms11257, doi: 10.1038/ncomms11257.

Narasimhan V, Danecek P, Scally A, Xue Y, Tyler-Smith C, Durbin R. BCFtools/RoH: a hidden Markov model approach for detecting autozygosity from next-generation sequencing data. Bioinformatics. 2016; 32(11):1749–1751. 10.1093/bioinformatics/btw044, doi: 10.1093/bioinformatics/btw044.

Ohm RA, Feau N, Henrissat B, Schoch CL, Horwitz BA, Barry KW, Condon BJ, Copeland AC, Dhillon B, Glaser F, Hesse CN, Kosti I, LaButti K, Lindquist EA, Lucas S, Salamov AA, Bradshaw RE, Ciuffetti L, Hamelin RC, Kema GHJ, et al. Diverse Lifestyles and Strategies of Plant Pathogenesis Encoded in the Genomes of Eighteen Dothideomycetes Fungi. PLOS Pathogens. 2012; 8(12):1–26. 10.1371/journal.ppat.1003037, doi: 10.1371/journal.ppat.1003037.

Okonechnikov K, Conesa A, García-Alcalde F. Qualimap 2: advanced multi-sample quality control for high-throughput sequencing data. Bioinformatics. 2016; 32(2):292–294. 10.1093/bioinformatics/btv566, doi: 10.1093/bioinformatics/btv566.

Oksanen J, Blanchet FG, Friendly M, Kindt R, Legendre P, McGlinn D, Minchin PR, O’Hara RB, Simpson GL, Solymos P, Stevens MHH, Szoecs E, Wagner H. vegan: Community Ecology Package; 2019, https://CRAN.R-project.org/package=vegan, r package version 2.5-6.

Regalado J, Lundberg DS, Deusch O, Kersten S, Karasov T, Poersch K, Shirsekar G, Weigel D. Combining whole-genome shotgun sequencing and rRNA gene amplicon analyses to improve detection of microbe–microbe interaction networks in plant leaves. The ISME Journal. 2020 Aug; 14(8):2116– 2130. https://www.nature.com/articles/s41396-020-0665-8, doi: 10.1038/s41396-020-0665-8, number: 8 Publisher: Nature Publishing Group.

Robinson RJ, Fraaije BA, Clark IM, Jackson RW, Hirsch PR, Mauchline TH. Endophytic bacterial community composition in wheat (Triticum aestivum) is determined by plant tissue type, developmental stage and soil nutrient availability. Plant and Soil. 2016; 405(1-2):381–396. 10.1007/s11104-015-2495-4, doi: 10.1007/s11104-015-2495-4.x

Saitou N, Nei M. The neighbor-joining method: a new method for reconstructing phylogenetic trees. Molecular Biology and Evolution. 1987 07; 4(4):406–425. 10.1093/oxfordjournals.molbev.a040454, doi: 10.1093/oxfordjournals.molbev.a040454.

Sartori M, Nesci A, García J, Passone MA, Montemarani A, Etcheverry M. Efficacy of epiphytic bacteria to prevent northern leaf blight caused by Exserohilum turcicum in maize. Revista Argentina de Microbiologia. 2017; 49(1):75–82. 10.1016/j.ram.2016.09.008, doi: 10.1016/j.ram.2016.09.008.

Sartori M, Nescia A, Formento Á, Etcheverry M. Selección de microorganismos epifíticos de maíz como potenciales agentes debiocontrolde Exserohilum turcicum. Revista Argentina de Microbiologia. 2015; 47(1):62–71. 10.1016/j.ram.2015.01.002, doi: 10.1016/j.ram.2015.01.002.

Savary S, Willocquet L, Pethybridge SJ, Esker P, McRoberts N, Nelson A. The global burden of pathogens and pests on major food crops. Nature Ecology & Evolution. 2019 Mar; 3(3):430–439. https://www.nature.com/articles/s41559-018-0793-y, doi: 10.1038/s41559-018-0793-y, number: 3 Publisher: Nature Publishing Group.

Schol-Schwarz MB. The genus Epicoccum Link. Transactions of the British Mycological Society. 1959; 42(2):149–IN3. 10.1016/S0007-1536(59)80024-3, doi: 10.1016/s0007-1536(59)80024-3.

Szabo LJ, Olivera PD, Wanyera R, Visser B, Jin Y. Development of a Diagnostic Assay for Differentiation Between Genetic Groups in Clades I, II, III, and IV of Puccinia graminis f. sp. tritici. Plant Disease. 2022 Aug; 106(8):2211–2220. https://apsjournals.apsnet.org/doi/10.1094/PDIS-10-21-2161-RE, doi: 10.1094/PDIS-10-21-2161-RE.

Trivedi P, Batista BD, Bazany KE, Singh BK. Plant–microbiome interactions under a changing world: responses, consequences and perspectives. New Phytologist. 2022; 234(6):1951–1959. https://nph.onlinelibrary.wiley.com/doi/abs/10.1111/nph.18016, doi: 10.1111/nph.18016.

Trivedi P, Leach JE, Tringe SG, Sa T, Singh BK. Plant–microbiome interactions: from community assembly to plant health. Nature Reviews Microbiology. 2020; 18(11):607–621. 10.1038/s41579-020-0412-1, doi: 10.1038/s41579-020-0412-1.

Van Overbeek L, Van Elsas JD. Effects of plant genotype and growth stage on the structure of bacterial communities associated with potato (Solanum tuberosum L.). FEMS Microbiology Ecology. 2008; 64(2):283–296. doi: 10.1111/j.1574-6941.2008.00469.x.

Vasimuddin M, Sanchit M, Li H, Aluru S. Efficient Architecture-Aware Acceleration of BWA-MEM for Multicore Systems. IEEE Parallel and Distributed Processing Symposium (IPDPS). 2019;.

Vidal-Villarejo M, Freund F, Hanekamp H, von Tiedemann A, Schmid K. Population genomic evidence for a repeated introduction and rapid expansion of the fungal maize pathogen Setospheria turcica in Europe. Genome Biology and Evolution. 2023 07; p. evad130. 10.1093/gbe/evad130, doi: 10.1093/gbe/evad130.

Ye SH, Siddle KJ, Park DJ, Sabeti PC. Benchmarking Metagenomics Tools for Taxonomic Classification. Cell. 2019; 178(4):779–794. 10.1016/j.cell.2019.07.010, doi: 10.1016/j.cell.2019.07.010.

Zhang J, Huang Y, Pu R, Gonzalez-Moreno P, Yuan L, Wu K, Huang W. Monitoring plant diseases and pests through remote sensing technology: A review. Computers and Electronics in Agriculture. 2019 Oct; 165:104943. https://www.sciencedirect.com/science/article/pii/S016816991930290X, doi: 10.1016/j.compag.2019.104943.

