## Supplemental File for "Regional Diversity and Leaf Microbiome Interactions of the Fungal Maize Pathogen *Exserohilum turcicum* in Switzerland: A Metagenomic Analysis"

#### – Supporting Information –

#### Contents

**Supplementary Figures** **2**

**Supplementary Tables** **9**

#### Supplementary Figures

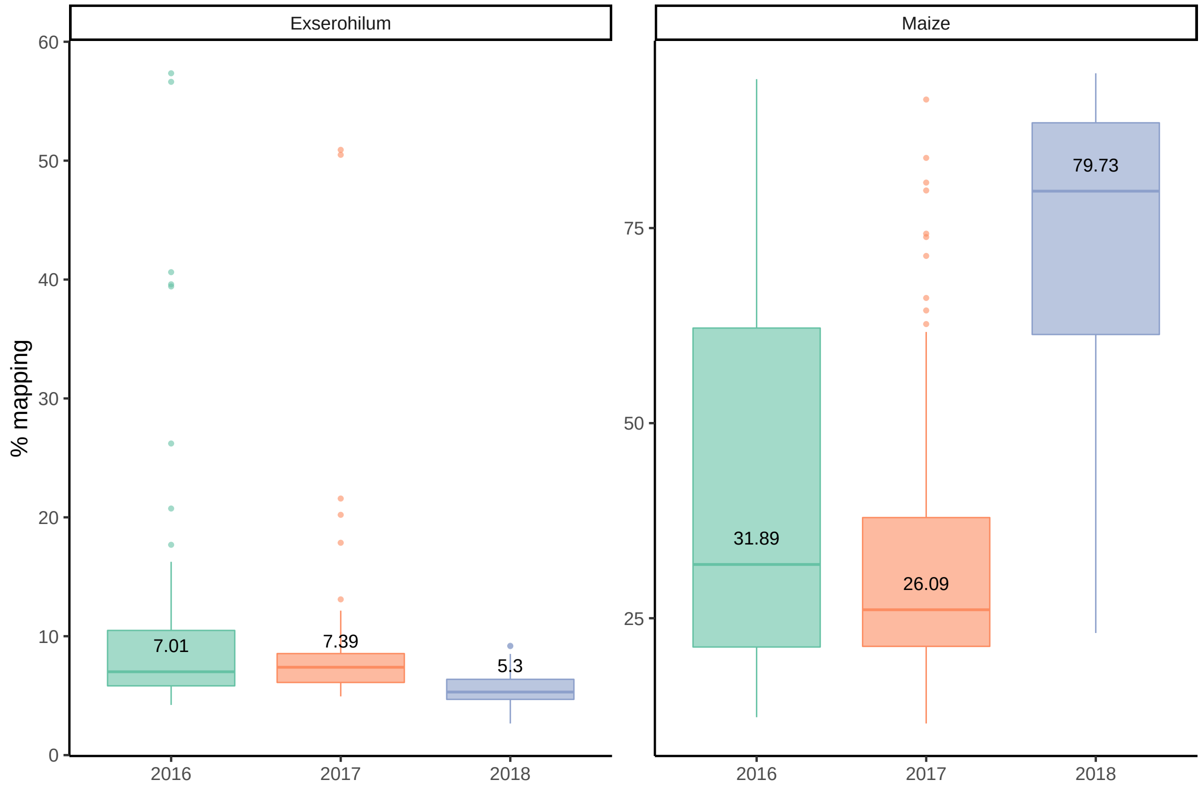


**Figure S1.** Summary of the percentage of reads mapping to *Exserohilum* and Maize reference genomes each year. Numbers within boxplots indicate the median.

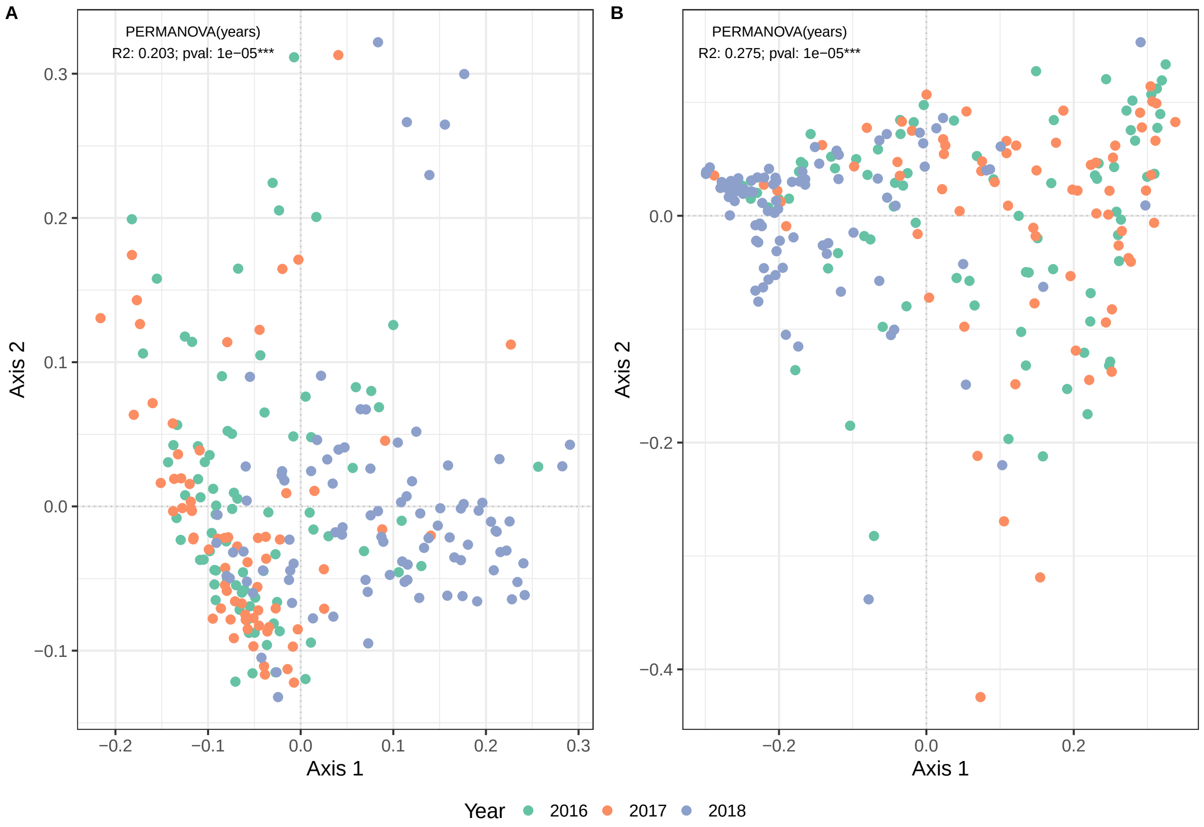


17

**Figure S2.** PCoA in **A)** Eukaryota and **B)** Bacteria separately with its corresponding $R^2$ and $p$ value of PERMANOVA analysis on the year.

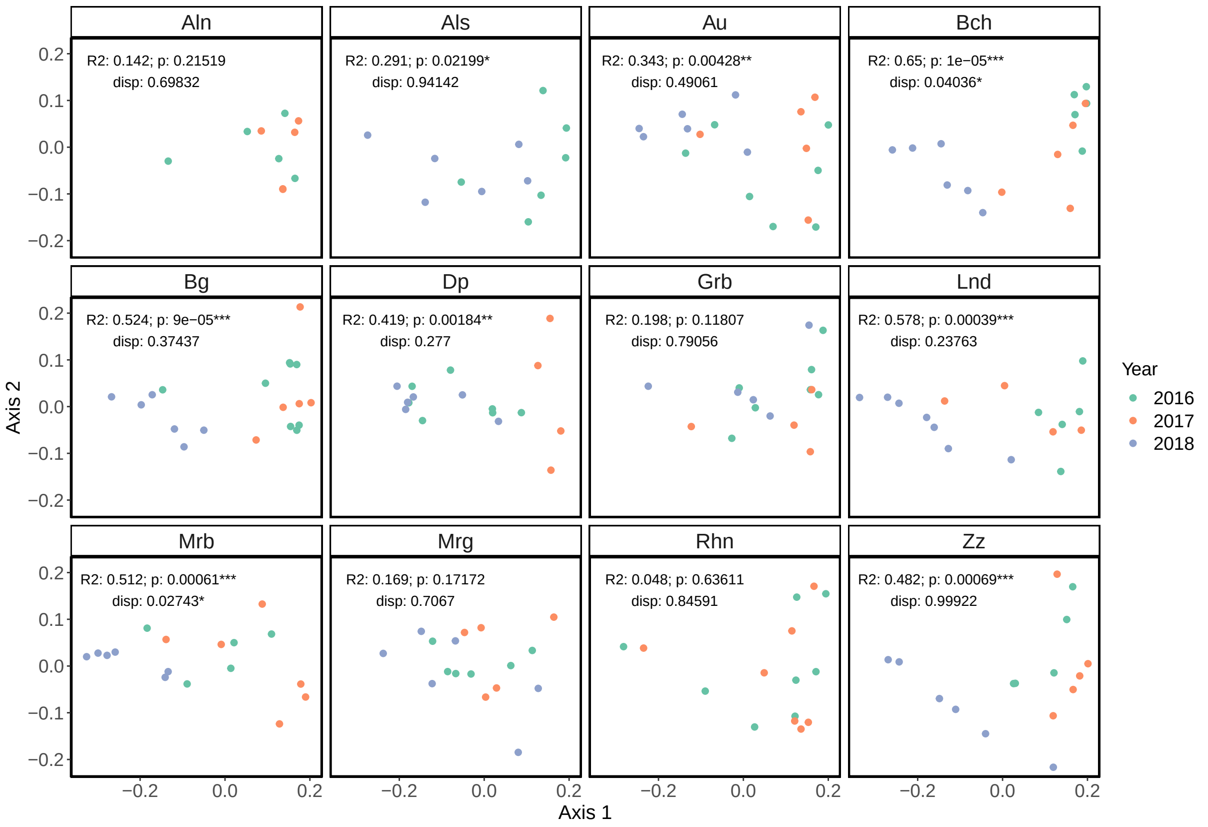


**Figure S3.** PCoA for each collection site sampled at least two years. $R^2$ and $p$-value of permanova analysis on the year and $p$ value of the test of homogenity of the dispersion (disp)

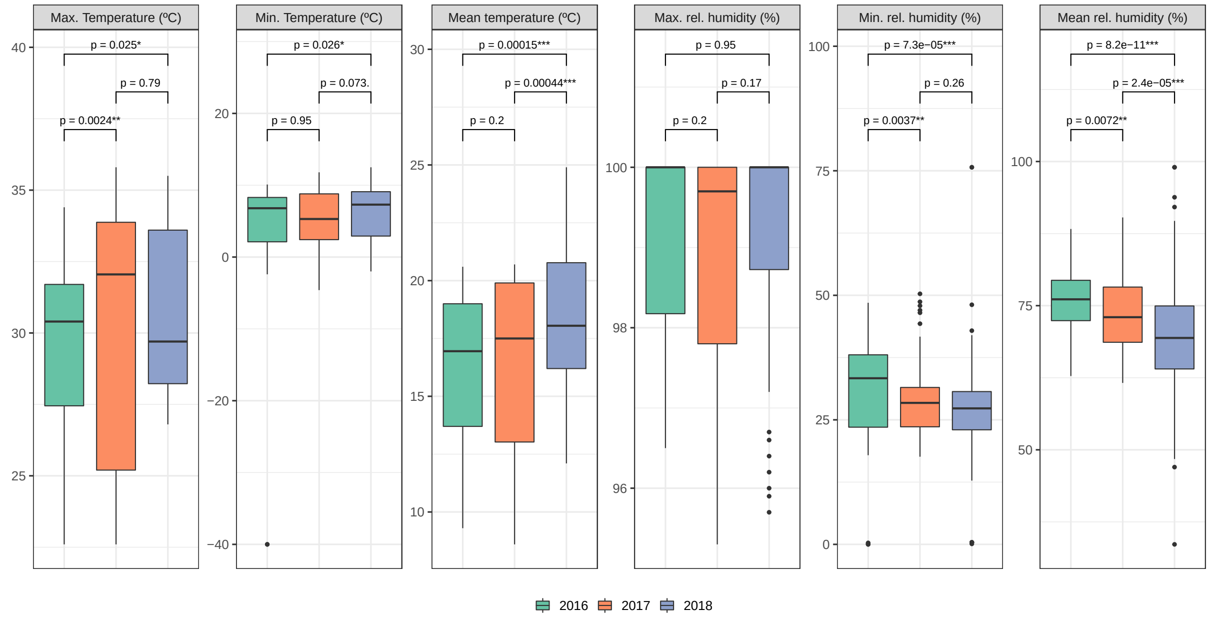


19

**Figure S4.** Minimum, maximum and mean temperature and relative humidity monthly records from Rheintal summarized by each year, including months from April to September. p indicates $p$ value of Wilcoxon rank test between years.

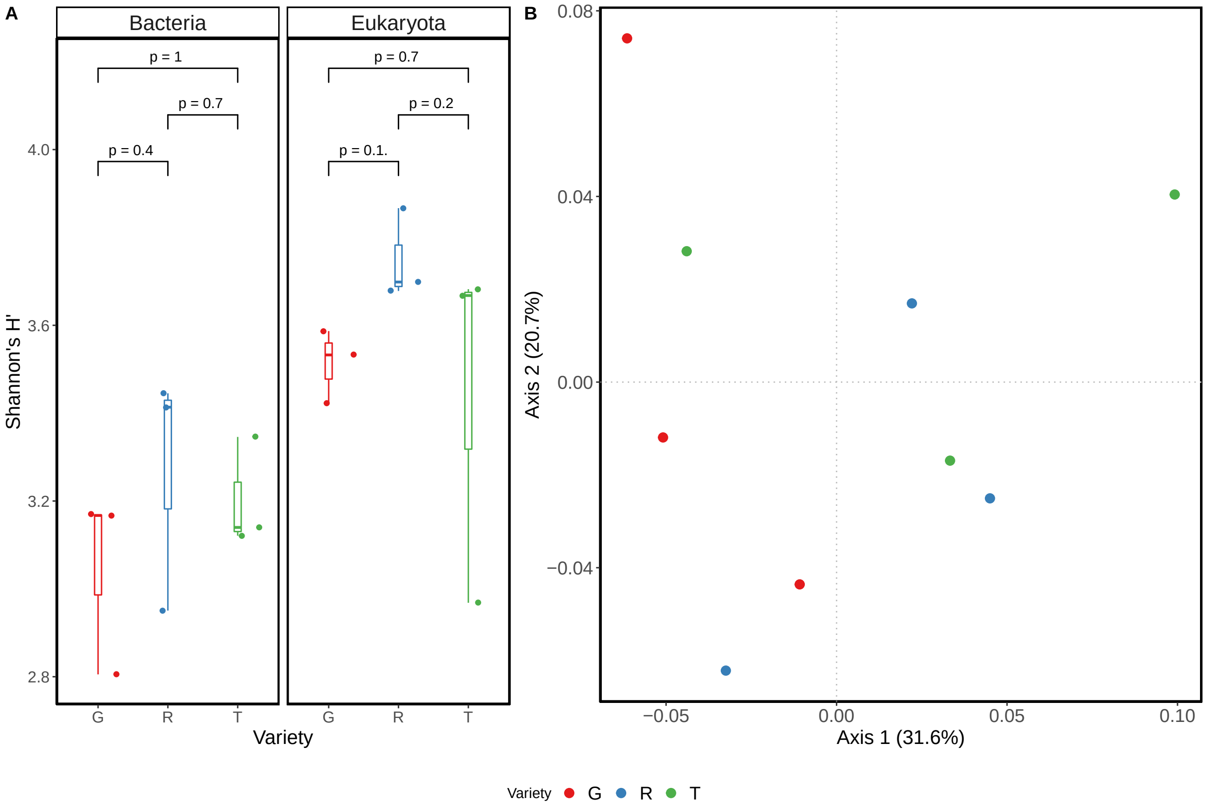


20

**Figure S5.** Pilot Study.**A)** Shannon's diversity index ($H'$) in bacterial and eukaryotic phyllobiome in the three maize varieties. p indicates $p$ value of Wilcoxon rank test between years. Size of 3 samples each variety. **B)** First two axis of the PCoA (and % of the variance) on the relative abundances of Eukaryota and Bacteria colored by the maize variety. PERMANOVA on the maize variety was not significant ($p = 0.1524$).

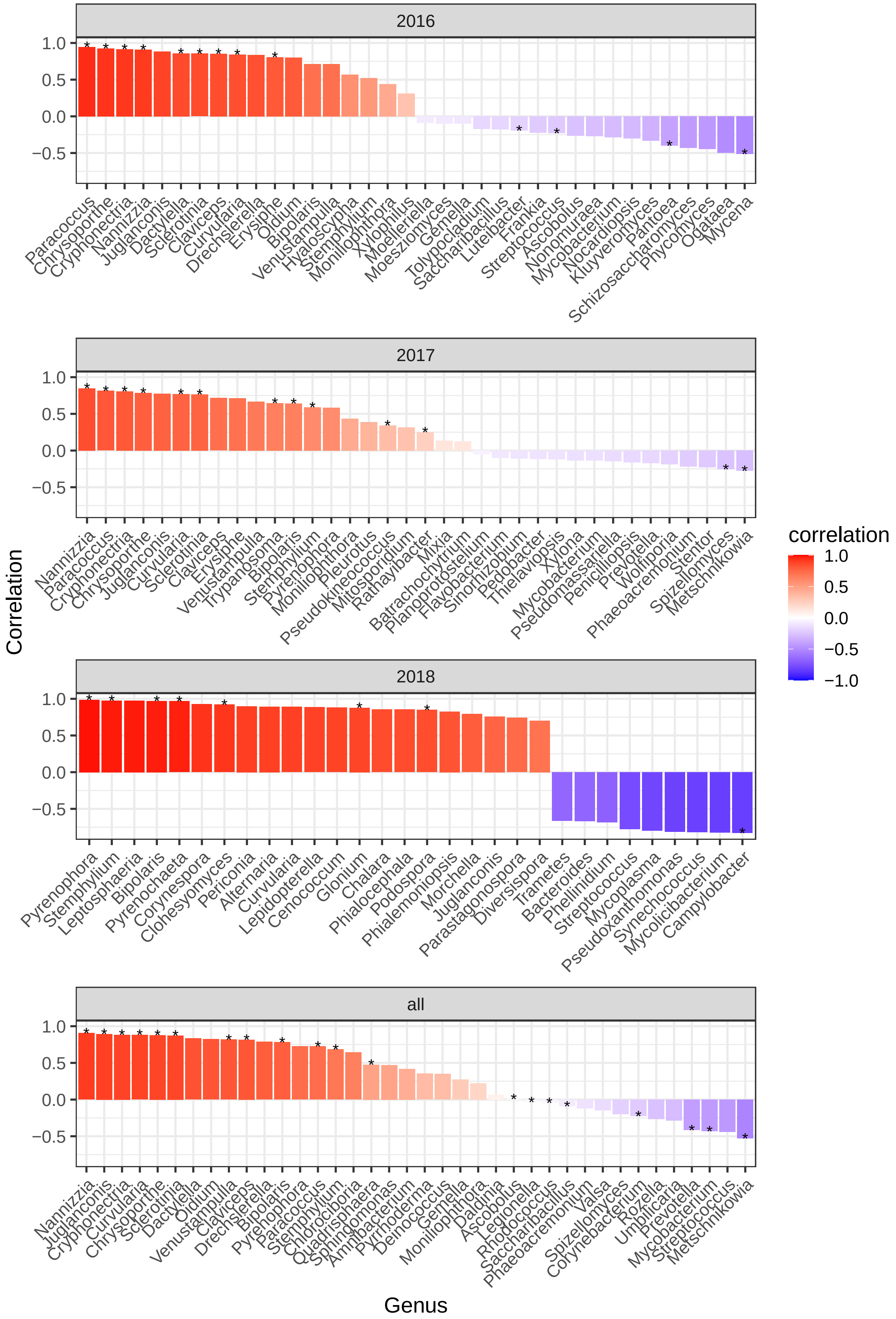


**Figure S6.** SparCC correlation coefficients with *Exserohilum* analyzing each separated and all together. Asterisk (*) denotes the interaction present in both the Neighborhood and Inverse Covarance methods of SPIEC-EASI

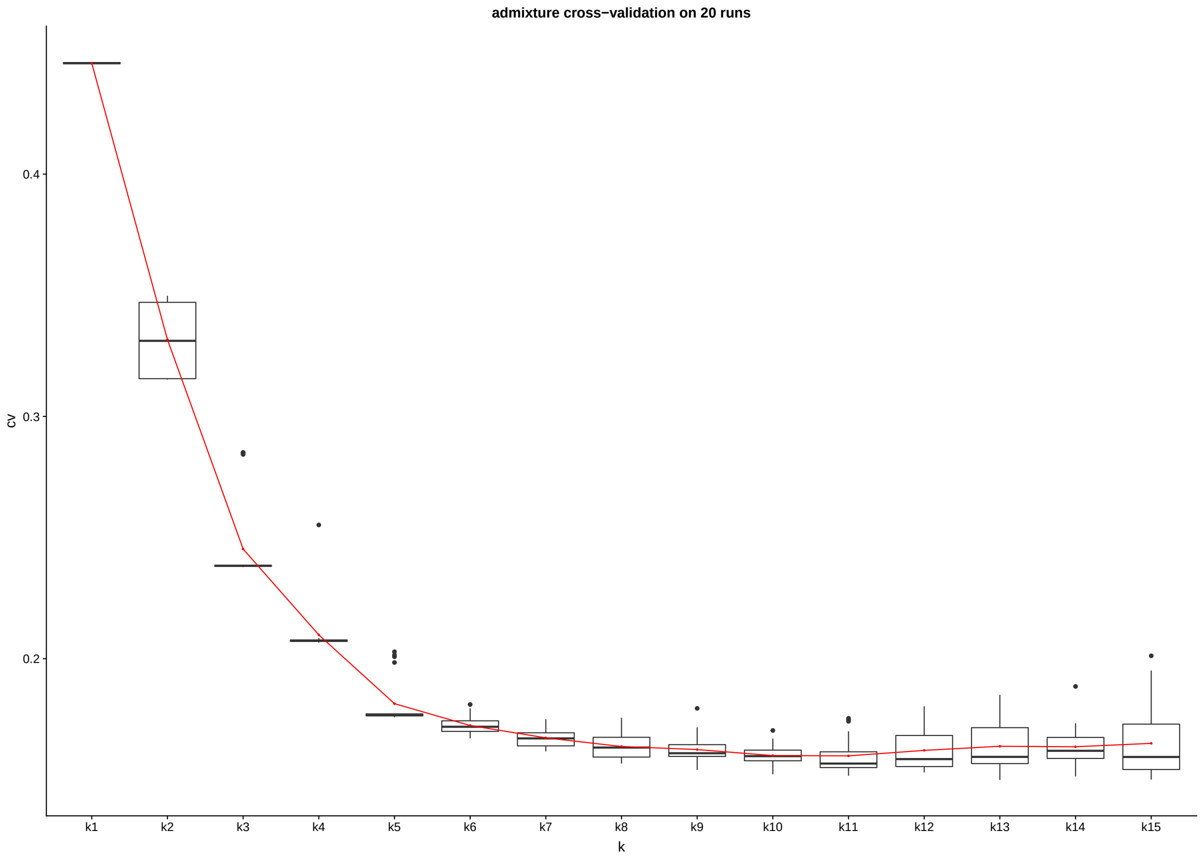


22

**Figure S7.** Cross-validation values on 20 independent runs in ADMIXTURE with K ranging from 1 to 15

### Supplementary Tables

**Table S1.** Multifactorial PERMANOVA test statistics for significance factors influencing phyllobiome dissimilarities and pairwise PERMANOVA on the year for Eukaryota and Bacteria separately.

| Source of variation | df | SS | MS | pseudo-F | $R^2$ | $p$ value |
|---|---|---|---|---|---|---|
| Year | | | | | | |
| *Eukarya* | 2 | 1.3877 | 0.6938 | 37.033 | 0.2030 | 1e-05 |
| *Bacteria* | 2 | 3.7989 | 1.8994 | 52.457 | 0.2755 | 1e-05 |
| Site | | | | | | |
| *Eukarya* | 20 | 0.9209 | 0.0460 | 2.457 | 0.1347 | 1e-05 |
| *Bacteria* | 20 | 1.4463 | 0.0723 | 1.997 | 0.1049 | 4e-04 |
| Latitude | | | | | | |
| *Eukarya* | 1 | 0.0203 | 0.0203 | 1.083 | 0.003 | 0.3407 |
| *Bacteria* | 1 | 0.1287 | 0.1287 | 3.554 | 0.0093 | 0.0296 |
| Longitude | | | | | | |
| *Eukarya* | 1 | 0.0689 | 0.0689 | 3.679 | 0.0101 | 0.0086 |
| *Bacteria* | 1 | 0.0584 | 0.0584 | 1.614 | 0.0042 | 0.1723 |
| Year x Site | | | | | | |
| *Eukarya* | 17 | 0.7084 | 0.0417 | 2.224 | 0.1036 | 2e-05 |
| *Bacteria* | 17 | 1.1513 | 0.0677 | 1.870 | 0.0835 | 0.0026 |
| Residuals | | | | | | |
| *Eukarya* | 199 | 3.7284 | 0.0187 | - | 0.5455 | - |
| *Bacteria* | 199 | 7.2056 | 0.0362 | - | 0.5227 | - |

df: degrees of freedom; SS: Sum of Squares; MS: Mean Squares
The analysis was performed with 99,999 permutations and calculated on Bray-Curtis dissimilarities.
Site refers to collection site.

**Table S2.** SparCC correlation by year (and all years togehter) with indication of the presence of direct interaction with *Exserohilum* defined in SPIEC-EASI. Table shows the top most negatively correlated genus with *Exserohilum* in all years ($< -0.2$ sparCC).

| | sparCC | | | | MB | | | | GL | | | |
|---|---|---|---|---|---|---|---|---|---|---|---|---|
| | All | 2016 | 2017 | 2018 | All | 2016 | 2017 | 2018 | All | 2016 | 2017 | 2018 |
| Metschnikowia | -0.529 | -0.422 | -0.274 | -0.624 | Yes | No | Yes | No | Yes | No | Yes | No |
| Streptococcus | -0.440 | -0.230 | 0.117 | -0.777 | Yes | Yes | No | No | No | Yes | No | Yes |
| Mycobacterium | -0.431 | -0.288 | -0.138 | -0.758 | Yes | No | Yes | No | Yes | Yes | No | No |
| Prevotella | -0.417 | 0.001 | -0.171 | -0.743 | Yes | No | No | No | Yes | No | Yes | No |
| Umbilicaria | -0.285 | 0.080 | 0.096 | -0.555 | Yes | No | No | No | No | No | No | No |
| Rozella | -0.266 | -0.040 | -0.183 | -0.330 | Yes | No | No | No | No | No | No | No |
| Corynebacterium | -0.224 | -0.149 | -0.078 | -0.578 | Yes | No | No | No | Yes | No | No | No |
| Spizellomyces | -0.201 | -0.321 | -0.255 | 0.031 | No | No | Yes | No | Yes | No | Yes | No |

MB: neighborhood selection; GL: inverse covariance selection
No: no direct interaction with *Exserohilum* in SPIEC-EASI; Yes: direct interaction with *Exserohilum* in SPIEC-EASI;
